# Eukaryotes’ closest relatives are internally simple syntrophic archaea

**DOI:** 10.1101/2025.02.26.640444

**Authors:** Hiroyuki Imachi, Masaru K. Nobu, Shun’ichi Ishii, Yuga Hirakata, Tetsuro Ikuta, Yuta Isaji, Makoto Miyata, Masayuki Miyazaki, Yuki Morono, Kazuyoshi Murata, Satoshi Nakagawa, Miyuki Ogawara, Satoshi Okada, Yumi Saito, Sanae Sakai, Shigeru Shimamura, Yuhei O. Tahara, Yoshihiro Takaki, Yoshinori Takano, Eiji Tasumi, Katsuyuki Uematsu, Toshihiro Yoshimura, Ken Takai

## Abstract

Eukaryotes are theorized to have originated from an archaeal phylum *Promethearchaeota* (formerly ‘Asgard’ archaea)^1,2^. The first cultured representatives revealed valuable insight^3,4^ but are distantly related to the first eukaryotic common ancestor (FECA), leaving many unknowns regarding this archaeon’s biology. Here, we report isolation of two strains belonging to the order proposed as FECA’s closest relative, ‘Hodarchaeales’,^5^ as members of a pure co- and tri-culture with methanogenic partners. Both are obligately anaerobic, syntrophic, peptide-degrading, and mesophilic archaea that have simple internal cell structure and produce protrusions and vesicles, like *Promethearchaeales*^3,4^. All strains demonstrate behavior focused on cell construction rather than division, unlike typical prokaryotes^6^. Strain HC1 possesses genes associated with aerobic lifestyles but lacks complete pathways for aerobic respiration and co-cultures cannot grow under (micro)aerobic conditions, suggesting the genes support oxygen detoxification rather than respiration. Reflecting this, HC1 can survive and grow under microaerobic conditions only when aerobic organisms are present. Phylogenetic analyses indicate FECA may have possessed these genes and thus some aerotolerance. Physiological, genomic, and phylogenetic observations indicate FECA was a simple-celled anaerobic syntrophic peptide-degrading archaeon with a non-growth-centric lifestyle and potential adaptations towards an oxygenated planet—the archaea-eukaryote transition was steep in both cell structure and aerobiosis.

## Introduction

Understanding how complex eukaryotic cells emerged from their prokaryotic ancestors is a fundamental challenge in biology. The widely accepted endosymbiotic theory posits that the first eukaryotic cells arose through the merging of an archaeal host cell and an alphaproteobacterial endosymbiont, which later became the mitochondria^7–10^. Although the identity of the host archaeon has long been enigmatic, recent metagenomic and phylogenetic analyses indicate that eukaryotes very likely evolved from within the archaeal group ‘Asgard’ archaea^5,11–13^, now formally classified as the phylum *Promethearchaeota* within the kingdom *Promethearchaeati*^2^. This relationship is further reflected in the presence of abundant eukaryotic signature proteins (ESPs) in their genomes^11,12,14^. In addition, several of these ESPs exhibit structural and functional similarities to those found in eukaryotes^15–18^. Recent studies have also reported that the antiviral defense systems in eukaryotes may have originated from this group of archaea^19,20^.

What remained unclear was the cell structure and physiology of the archaeal group, including FECA, with both having significant implications for how archaea may have transitioned towards eukaryotes. Based on the presence of abundant and diverse ESPs in these archaea, studies hypothesized that FECA may have already had complex internal cell structure (*e.g.*, endomembrane systems) prior to endosymbiosis of the alphaproteobacterial ancestor of mitochondria (that is, “mito-intermediate” or “mito-late”)^1,21,22^. Cultivation efforts yielded one isolate *Promethearchaeum syntrophicum* strain MK-D1^1,3^ and an enrichment culture predominated by *Candidatus* (*Ca.*) ‘Lokiarchaeum ossiferum’^4^, both revealing typical prokaryotic internal structure^3,4^ and potential evidence for an actin-based cytoskeleton^4,23^. Both mesophilic strains gain energy from anaerobic degradation of peptides/amino acids through symbiotic metabolic (that is, syntrophic) interactions with H_2_-consuming methanogenic archaea or sulfate-reducing bacteria^2–4^. Metagenomics studies have also speculated that H_2_-mediated syntrophy may have played an important role, though with a different partner^21^. Collectively, these pointed towards an anaerobic syntrophic FECA with simple cell structure and a steep transition in physiology and structure between archaea and eukaryotes.

However, studies have presented doubts on whether features found in these *Promethearchaeia* (formerly ‘Lokiarchaeia’) cultures are relevant to FECA^1,5,23^. First, metagenomic predictions suggest a wide diversity of physiologies across *Promethearchaeota* (*e.g.*, autotrophy, organotrophy, and rhodopsin-based phototrophy)^21,24–26^. Second, recent phylogenetic analyses identify other lineages of *Promethearchaeota*, *Ca.* ‘Heimdallarchaeia’ or its order *Ca.* ‘Hodarchaeales’, that branch closer to eukaryotes^5,13^. Based on these observations, a variety of features have been proposed for FECA, including complex internal cell structure^3,5,9,21,22,27^. One potential feature of FECA that has also recently attracted attention is aerobic respiration^5,21,14,26,27^. Studies find that, unlike *Promethearchaeia*, some ‘Heimdallarchaeia’ possess terminal oxidases^5,21^ and further propose FECA may have been aerobic^14,26,27^. Collectively, these suggest that FECA already had the beginnings of structural and respiratory features associated with eukaryotes. However, the cell structure of ‘Heimdallarchaeia’ has yet to be elucidated. Moreover, as many obligately anaerobic organisms possess terminal oxidases for oxygen tolerance^28,29^, the simple presence of such enzymes cannot tell us whether an organism is an aerotolerant anaerobe (O_2_-depletion hypothesis^30^) or (facultative) aerobe. In light of this, cultivation of class ‘Heimdallarchaeia’ and its order ‘Hodarchaeales’ is essential to unravel the cell structure and physiology of FECA^1^.

In this study, we successfully cultured two strains of ‘Hodarchaeales’, one isolated in a pure syntrophic co-culture and another in a tri-culture with methanogenic partners. We further report the cultivation procedures, cellular structures, physiological characteristics, and genomic features of these strains, and their implications for the origin of eukaryotes.

### Cultivation of novel ‘Hodarchaeales’ strains

To culture members of ‘Heimdallarchaeia’ and explore their potential relationship with oxygen, we capitalized on previous experience in cultivation of *P. syntrophicum* MK-D1^3,31^. We attributed successful isolation of this strain to a long-term bioreactor-based increase of metabolic activity, acclimation to artificial conditions, and selection against other populations (that is, enrichment)^2,3,31^. Previous studies also hint that the presence and activity of anaerobic methane-oxidizing microbial consortia were likely important in supporting the growth of *Promethearchaeota* members, presumably through the excretion of organic compounds^31,32^. Here, we looked to a gas production well in the Boso Peninsula, Chiba Prefecture, Japan, which is operationally similar to our previous cultivation methane-fed “bioreactor” (Extended Data Fig. 1). Methane-bearing ancient seawater (*i.e*., brine) upwells into the well and supports a biofilm on the walls containing populations identical or similar to those found in marine sediments, according to 16S rRNA gene tag-sequencing (iTag) analysis^33^. Unlike our previous anoxic reactor, the biofilm is intermittently exposed to air (and thus oxygen) due to periodic eruptions of brine, akin to geysers, that cause the water level to drop temporarily^33^. Enriched in these biofilms were small populations of ‘Heimdallarchaeia’, which we targeted for isolation (Supplementary Table 1).

Aiming to culture the biofilm microbial community in the laboratory while mimicking natural conditions, we used filtered brine from the well as the basal medium and supplemented this with methane and magnetite to attempt “co-enrichment” of *Promethearchaeota* members with anaerobic methane-oxidizing (ANME) archaea (Supplementary Note 1). After approximately three months, we detected several 16S rRNA gene sequences of ‘Hodarchaeales’, ‘Heimdallarchaeales’, and *Promethearchaeales* (Fig. 1a and Supplementary Table 1). We implemented proteinaceous substrates, casamino acids and powdered milk (each 0.005%, w/v), as energy sources for sub-cultures to enhance growth of these archaea, based on our previous observation that syntrophic degradation of amino acids/peptides as a widespread feature across the phylum^3^ (Supplementary Note 1 and Supplementary Table 1). To also select against other organisms, we supplemented the medium with five antibiotics: ampicillin, vancomycin, kanamycin, streptomycin, and erythromycin.

**Fig. 1.**
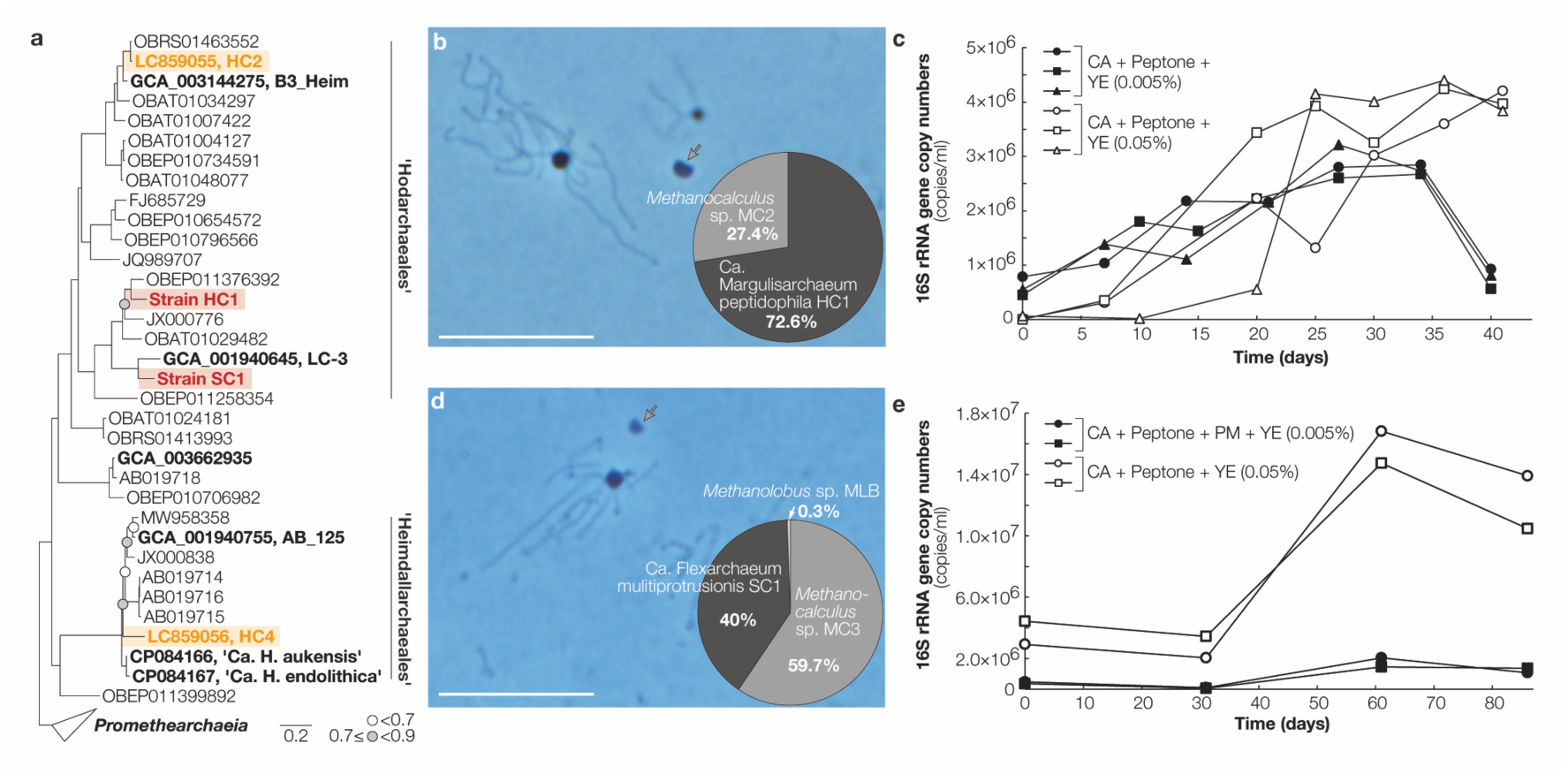
16S rRNA gene-based phylogeny, photomicrographs and growth curves of cultured ‘Hodarchaeales’ strains HC1 and SC1. **a**, Maximum-likelihood estimation of the phylogeny of 16S rRNA gene sequences, showing the phylogenetic position of cultured strains. The 16S rRNA genes obtained from isolated strains, enrichment culture, and metagenome assemblies are shown in red, orange, and black bold, respectively. Bootstrap values around critical branching points are also shown. **b**, Phase-contrast photomicrograph of pure co-culture of HC1 and *Methanocalculus* sp. MC2. The relative abundance of each member (in late exponential phase) was estimated using 16S rRNA gene-tag-sequencing (iTag) analysis and shown as a pie chart. **c**, Growth curves of HC1 in anaerobic medium supplemented with casamino acids (CA), peptone and yeast extract (YE) (0.005% w/v each). **d**, Phase-contrast photomicrograph of strain SC1 in a tri-culture with a pie chart indicating the relative abundance of each member in late exponential phase based on iTag analysis. **e**, Growth curves of SC1 in anaerobic medium supplemented with CA, peptone, powdered milk (PM) and YE (0.005%, 0.005%, 0.005%, and 0.05% w/v, respectively). Gray arrows indicate *Methanocalculus* cells. Scale bars, 10 µm.

For cultures that sustained *Promethearchaeota* members according to iTag analysis, we sub-cultured twice in fresh medium of the same composition at roughly 100-day intervals and, for cultures that enriched one population of ‘Hodarchaeales’ over other *Promethearchaeota*, further sub-cultured using a synthetic medium mimicking the natural brine and supplemented with casamino acids, powdered milk, and five antibiotics (Fig. 1a and Supplementary Table 1). To further simulate the natural conditions, we omitted reducing agents (sodium sulfide and cysteine) to leave trace amounts of oxygen in the medium and select for ‘Hodarchaeales’ over *Promethearchaeales* members, each presumably associated with more “oxidized” and “reduced” lifestyles^2,21^. After four successive transfers under the same cultivation conditions, while aerobic bacteria (that is, *Mariniphaga* and *Alcanivorax*) dominated the culture, the ‘Hodarchaeales’ population increased to 1.8% in relative abundance and *Promethearchaeales* members were no longer detectable (Supplementary Table 1). This suggested that the ‘Hodarchaeales’ population may have weak aerotolerance and depend on oxygen consumption by the aerobic bacteria.

In subsequent sub-cultures, reducing agents were re-introduced and peptone was added as an additional energy source. The ‘Hodarchaeales’ population reached up to 30% in relative abundance and maximum cell densities up to 10^6^ 16S rRNA gene copy numbers ml^-1^. As for the other populations, only *Mariniphaga*, *Alcanivorax* and *Methanocalculus* remained (Supplementary Table 1). We isolated the *Mariniphaga* and *Alcanivorax* species, identified antibiotics they are sensitive to (Supplementary Table 2), and applied these antibiotics (gentamycin, ciprofloxacin, cephalothin, or ofloxacin) to the ‘Hodarchaeales’-dominated cultures (Supplementary Table 1). In parallel we also optimized the cultivation temperature and energy source types and concentrations (Extended Data Fig. 2 and Supplementary Table 1). Three years after the original sampling, we isolated ‘Hodarchaeales’ in a purified co-culture with a single partnering methanogen population (*Methanocalculus*), each designated as strains HC1 and MC2, in an anaerobic medium supplemented with casamino acids, peptone and yeast extract (each 0.005%, w/v) at 30°C (Fig. 1 and Supplementary Table 1). Under this cultivation condition, HC1 required 25–35 days to reach full growth with a doubling time of approximately 7–12 days, reaching a maximum cell density of 10^6^ 16S rRNA gene copies ml^-1^ (Fig. 1c). HC1 cells were morphologically similar to *Promethearchaeales* members^2–4^, with coccoid cells, albeit with larger diameters of 0.7–1.8 µm (1.3 µm on average; Supplementary Table 3), and multiple protrusions extending from the cell surface (Fig. 1b).

Using a cultivation strategy similar to that for HC1, we successfully isolated another ‘Hodarchaeales’ strain and also enriched other ‘Hodarchaeales’ and ‘Heimdallarchaeaceae’ members (Fig. 1, Supplementary Table 1, and Supplementary Text 2). One ‘Hodarchaeales’ strain, designated as SC1, was isolated in a purified tri-culture with *Methanocalculus* sp. strain MC3 and *Methanolobus* sp. strain MLB, with relative abundances of approximately 40%, 60%, and <0.3%, respectively, based on iTag analysis (Fig. 1d and Supplementary Table 1). The tri-culture is stably maintained in anaerobic medium supplemented with casamino acids, powdered milk, polypeptone peptone (0.01%, w/v), and yeast extract (0.05%, w/v). SC1 required approximately 60 days to reach maximum growth with a doubling time of approximately 14–20 days, reaching a maximum cell density of 10^7^ 16S rRNA gene copies (Fig. 1e). The 16S rRNA gene sequence identity between SC1 and HC1 was 77.75% based on BLAST analysis. SC1 cells were morphologically similar to HC1 but with a smaller diameter, 0.6–1.2 µm (0.9 µm on average) (Fig. 1d).

We sequenced the genomes of both strains. HC1 and SC1 possess circular genomes 7.91 Mb and 6.75 Mb, respectively (Supplementary Tables 4‒6). HC1 has a larger genome than all reported ‘Heimdallarchaeia’ metagenome-assembled genomes^27^. Phylogenetic analyses showed that both strains belonged to the family-level clade LC-3 in ‘Hodarchaeales’ (Fig. 1a). We tentatively propose the names *Ca*. ‘Margulisarchaeum peptidophila’ and *Ca.* ‘Flexarchaeum multiprotrusionis’ for strains HC1 and SC1, respectively, under the family *Ca*. ‘Margulisarchaeaceae’ (Supplementary Note 3).

### Cells lack internal complexity

Although several studies speculate that *Promethearchaeota* more closely related to eukaryotes may have complex cell structure distinct from *P. syntrophicum* MK-D1^1,21,22^, microscopic observation of HC1 and SC1 cells revealed that both have signatures of typical prokaryotic cells, including simple internal structure (Fig. 2 and Extended Data Fig. 3). Cells are coccoid and generally found as individuals, occasionally in aggregates (Fig. 2a, o, y and Extended Data Fig. 3c, p). Observation of the internal structure of HC1 and SC1 using transmission electron microscopy (TEM) and cryo-EM revealed no endomembrane systems (Fig. 2g, h, k–n, u, v, y–ab, Supplementary Video 1). We infrequently observed cells with low-electron density intracellular vesicle-/vacuole-like structures (Fig. 2m, n, and ab). While a previous study proposed *Promethearchaeota* may have membrane-based compartmentalization of the chromosome based on the observation of spatially localized genomic material^34^, we show that no membrane structures are involved in this segregation using fluorescence in situ hybridization (FISH), TEM, and correlative light-electron microscopy (CLEM) (Fig. 2, Extended Data Figs. 3 and 4, Supplementary Fig. 1, and Supplementary Video 2).

**Fig. 2.**
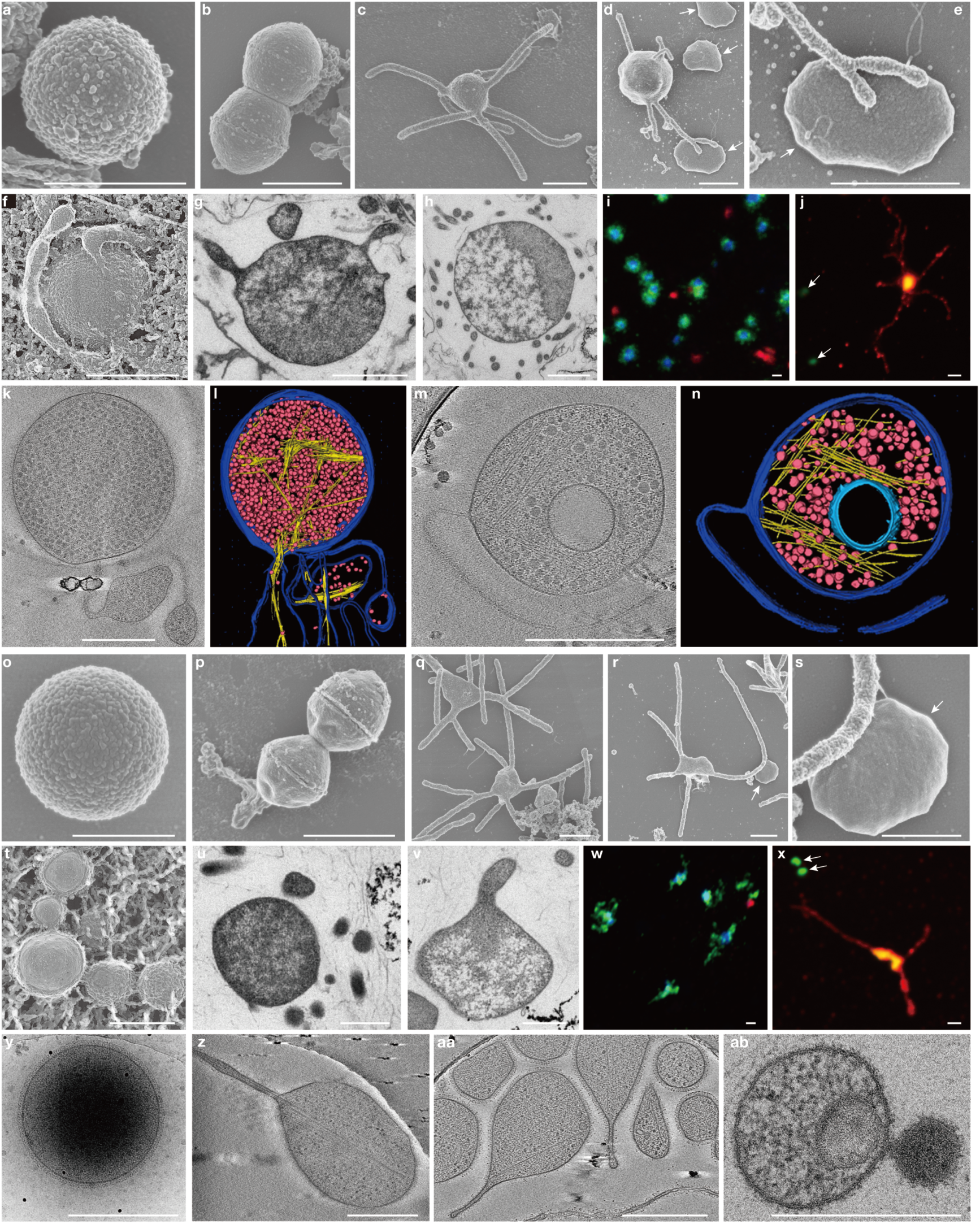
Microscopy characterization of HC1 and SC1. **a–n**, HC1 cultures. **a**–**e**, SEM images showing single (**a**), dividing (**b**), protrusion-producing (**c**), and methanogen-attached (**d**) cells and a magnified view of **d** (**e**). **f**, QFDE image. **g**, **h**, Ultrathin section. **i**, Fluorescence image of cells stained with DAPI (blue) and hybridized with nucleotide probes targeting HC1 (green) and *Methanocalculus* (red). **j**, Immunofluorescence image of cells stained with SYBR Green I (yellow/green) and labeled with an HC1 actin-specific antibody (red) (Extended Data Fig. 7). **k**–**n**, Slice images from cryo-tomograms (**k**, **m**) and corresponding 3D volume segmentations (**l**, **n**). Vacuole- or intracellular vesicle-like structures were infrequently observed (**m**, **n**). **o–ab**, SC1 cultures. **o**–**s** SEM images showing single (**o**), dividing (**p**), protrusion-producing (**q**), and methanogen-attached (**r**) cells and a magnified view of **r** (**s**). **t**, QFDE image. **u**, **v**, Ultrathin section. **w**, Fluorescence image of cells stained with DAPI (blue) and hybridized with nucleotide probes targeting SC1 (green) and *Methanocalculus* (red). **x**, Immunofluorescence image of cells stained with SYBR Green I (yellow/green) and labeled with an SC1 actin-specific antibody (red) (Extended Data Fig. 7). **y**, Cryo-EM snapshot. **z**, **aa**, Slice images from cryo-tomograms. **ab,** Ultrathin section of a cell containing a vacuole- or intracellular vesicle-like structure. White arrows indicate *Methanocalculus*. Images **c**, **f**, **g**–**n**, **t**–**x**, **z**– **ab** were obtained from cultures in late-exponential phase and others from mid-exponential phase. SEM, QFDE, ultrathin section, FISH, and immunofluorescence images are representative of *n*=98, 48, 60, 21, and 36 recorded images for HC1 and *n*=46, 44, 14, 9, and 11 recorded images for SC1. For HC1 and SC1, cryo-tomography images were obtained from 8 and 5 recorded tomograms, respectively. Scale bars, 1 µm (**a**–**j**, **p**– **r**, **w**, **x**) and 500 nm (**k**, **m**, **o**, **s**–**v**, **y**–**ab**).

Cells divide through binary fission and, during this, exhibit a ring-like structure (Fig. 2b, p). Both strains produced membrane vesicles (Extended Data Fig. 3i–m, z). Under cryo-EM and quick-freeze deep-etch replica electron microscopy (QFDE-EM), we could observe regular S-layer-like structures for SC1, which are typical of archaea^35^ (Fig. 2f, t, y, aa, and Extended Data Fig. 3g, z). A combination of protein multimer structure prediction via AlphaFold3^36^ and glycoproteomics revealed S-layer proteins conserved across many *Promethearchaeota* lineages that form symmetric homotetramers with a central vertical pore and are *N-*glycosylated like S-layer proteins of other archaea^35^ (Extended Data Fig. 5 and Supplementary Table 7).

After mid-exponential phase, most cells possess multiple membrane-based cytosol-connected protrusions of uniform diameter (roughly 340 nm), varying lengths, branching structure, and bulbous expansions (Fig. 2, Extended Data Fig. 3), morphologically similar to *P. syntrophicum* MK-D1^2,3^ and ‘L. ossiferum’^4^. Protrusions were not observed in dividing cells, suggesting that protrusion generation and cell division are mutually exclusive processes. The protrusion volume per cell was roughly 1.3 µm^3^ or 108% of the main cell body volume (Supplementary Table 3). Notably, both protrusions and cells were flexible and observed to sway by Brownian motion (Supplementary Videos 3 and 4). Fixation with formaldehyde and glutaraldehyde shrunk cells and formaldehyde tended to also disconnect their protrusions (Extended Data Fig. 6). As also observed for ‘L. ossiferum’^4^, actin filament-like structures were present in both strains and often extended from the cell interior towards the protrusions (Fig. 2j–n, x, z, aa, Extended Data Figs. 3h, y, z and 7, and Supplementary Fig. 2). Given the morphological similarity between all *Promethearchaeota* cultures, we can deduce that FECA likely had a similar cell structure and the protrusions play an important role in *Promethearchaeota* physiology, especially given their large volume.

As one major difference from the protrusions of *Promethearchaeales* (*P. syntrophicum* MK-D1^2,3^ and ‘L. ossiferum’^4^), the ‘Hodarchaeales’ protrusions were discovered to have filaments on their surfaces (Fig. 2d, e, q–s, and Extended Data Fig. 3e, f, r, s). These filaments were helical, often short but variable in length (∼125 nm and ∼1.5 µm for HC1 and SC1), thicker than pili and archaella but comparable to flagella (15–20 nm and 20–25 nm in diameter for HC1 and SC1), and equipped with round structures at the tip (approximately 50 nm in diameter for HC1 and 55 nm for SC1) (Extended Data Fig. 3f, u–w). HC1 and SC1 cells are often observed attached to their partner methanogens via their protrusions, and these filaments tend to be present at the interface (Fig. 2d, e, r, s and Extended Data Fig. 3e, f, r, s). While the nature or function of these filaments remain unknown, these observations suggest that they may help mediate physical attachment and symbiotic interactions with partner organisms. Notably, prokaryotic filament structures—flagella, pili, and archaella—have been repeatedly demonstrated to play critical roles in syntrophic interactions (that is, partner recognition and electron transfer)^37–39^.

Intracellular granular structures were found in HC1 and SC1, albeit less abundant (Fig. 2k–n, z, aa and Extended Data Fig. 3h, x–z, Supplementary Video 1). A wide range of diameters, 11.7 to 46.1 nm (average of 24.8 ± 5.3 nm), was observed for these granules (Supplementary Fig. 3). In the transcriptome and proteome, we find high expression of two proteins that can form granules in this size range: ribosomes and ferritin(-like) proteins^40^. Among these, ferritin is detected among the top three highest expressed proteins in the transcriptome, at a gene expression level 1.77 times that of the sum of all ribosomal proteins (Supplementary Table 5).

### Strict dependency on syntrophy and peptides

The HC1 co-culture and SC1 tri-culture both contain H_2_-utilizing methanogenic archaea and addition of a methanogenesis inhibitor, 2-bromoethanesulfonate (10 mM), to the cultures completely suppressed growth. This shows that both HC1 and SC1 depend on H_2_-mediated symbiosis with methanogens. Such a metabolic symbiosis is known as syntrophy, in which an orgranotroph performs a hydrogen and/or formate-yielding metabolism that is thermodynamically inhibited by these products and a partner organism scavenges these products to low non-inhibitory concentrations^41^. Accordingly, amending high concentrations of hydrogen gas or formate also suppressed the growth of both strains (Supplementary Table 8). In the SC1 tri-culture, we detected a methanogen that utilizes only methylated compounds, *Methanolobus* sp. MLB, and suspect that SC1 may generate such compounds during peptide degradation (*e.g.*, methylsulfides from methionine degradation; Supplementary Table 6). Given that syntrophy and symbiosis have been hypothesized to be relevant to the origin of eukaryotes^3,7,9,21^, the discovery these features in ‘Hodarchaeales’ has major evolutionary implications (see “Implications for eukaryogenesis”).

Both strains are clearly dependent on peptides as supplementation with non-proteinaceous substrates did not stimulate growth of HC1 and SC1, and neither strain could grow when peptide substrates were replaced with a mixture of 20 proteinogenic amino acids (Fig. 3a and Supplementary Table 8). Ultra-performance liquid chromatography (UPLC)-based analysis of macromolecule distributions in the cultures indicated clear consumption of higher molecular weight molecules (Fig. 3b, c, Supplementary Fig. 4, Supplementary Note 4). Genome analyses revealed both strains encode extracellular peptidases, peptide transporters, amino acid degradation pathways of glutamate, aspartate, threonine, methionine, glycine, serine, alanine, and cysteine in both strains, and NiFe hydrogenases for syntrophic peptide degradation (Supplementary Tables 5 and 6). Transcriptomic and proteomic data further confirmed their expression (Supplementary Tables 5 and 6). Glycoproteomics revealed that peptide/nickel transport system substrate-binding proteins of both strains were highly *N-*glycosylated like *P. syntrophicum* MK-D1^42^ (Extended Data Fig. 8, Supplementary Table 7, and Supplementary Note 5), further evidencing the importance of peptides for the growth of these strains. Collectively, we demonstrate that both strains are obligately syntrophy-dependent peptidotrophic archaea. Notably, among the potential substrates tested, pyruvate suppressed growth at 1 mM but not at 100 µM (Supplementary Table 8). While the mechanism remains unclear, this sensitivity to pyruvate is an interesting parallel to the mitochondrion’s ability to oxidize pyruvate.

**Fig. 3.**
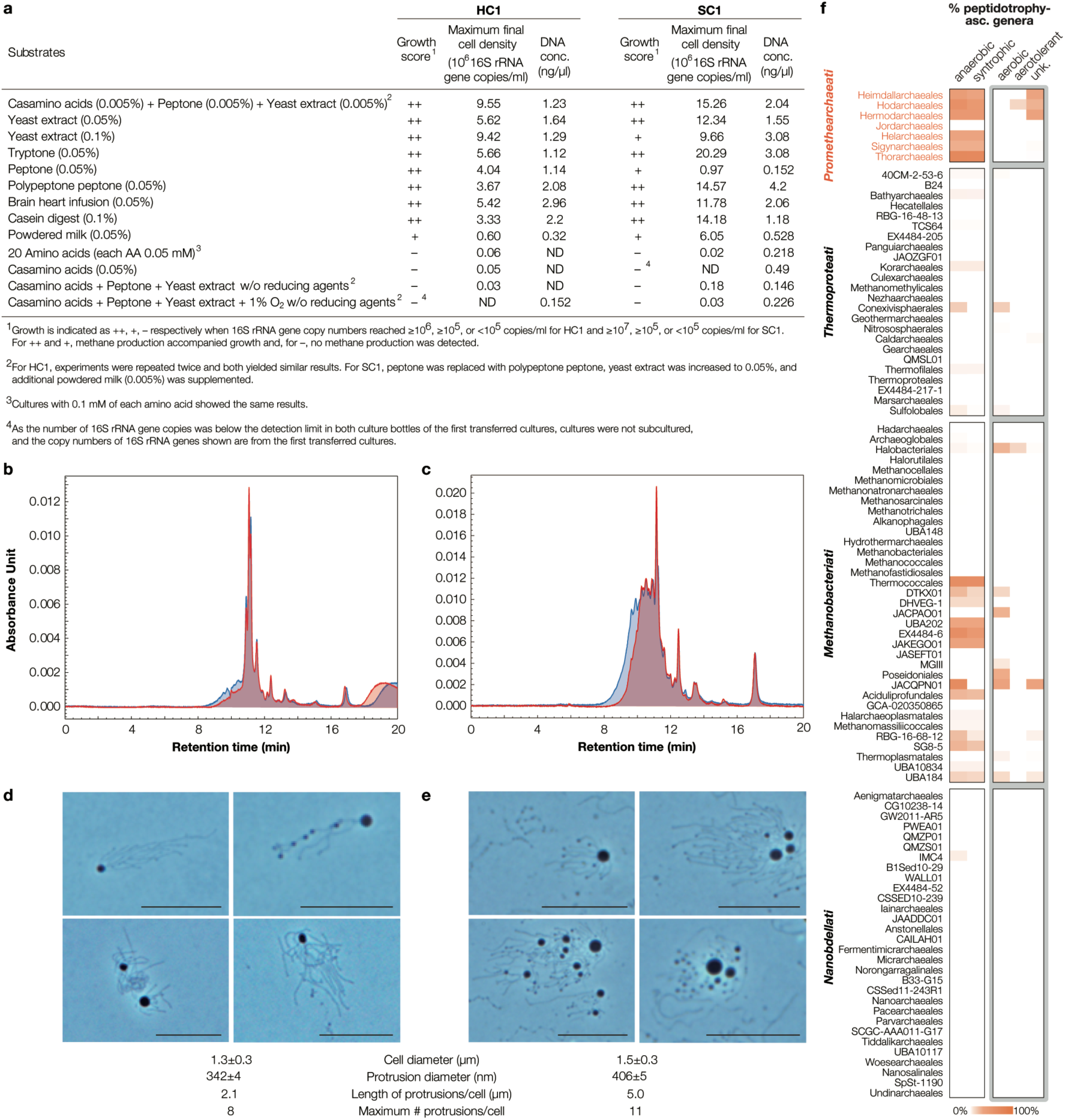
Peptide utilization by HC1 and SC1. **a,** Growth of HC1 and SC1 in cultures amended with complex peptide-containing organic carbon/energy sources and/or amino acids. For each cultivation condition, duplicate cultures were subcultured twice and quantification of growth based on 16S rRNA gene copies and DNA concentration are shown for the second subculture. Methane production was detectable in all cultures that showed positive growth (+ or ++). For HC1 and SC1, samples were taken 25 and 52 days after inoculation, respectively. The DNA concentration is based on DNA extracted from 1 mL of culture medium dissolved in 21 µl tris(hydroxymethyl)aminomethane-ethylenediaminetetraacetic acid (TE) buffer. **b, c,** UPLC-based detection of high molecular weight compounds in HC1 cultures fed with (**b**) casamino acids, peptone, and yeast extract and (**c**) peptone. **d, e,** Phase-contrast microscopy images of HC1 with (**d**) 0.005% and (**e**) 0.05% w/v yeast extract. The corresponding average cell diameter of unfixed cells, protrusion diameter for unfixed cells, estimated protrusion length of fixed cells, and maximum number of protrusions observed per cell (also see Supplementary Table 3). The scale bars are 10 µm. **f,** Capacities to perform anaerobic, syntrophic, aerobic, and aerotolerant peptidotrophy predicted for archaea based on gene expressivity (see Methods; Supplementary Table 12). Archaea that (i) possess a low-expressivity Cox or Cbd or (ii) a high-expressivity cox but lacking complex III were annotated to have an unknown O_2_ utilization capacity (“unk.”). Within each genus, the percentage of species with the above abilities was calculated and the average is shown for each order. Orders containing less than three genera are not shown (see Supplementary Table 12).

In cultures of HC1, we observed that an increasing amount of peptides added to the medium increased the cell size and protrusion number but not the growth rate, a trend atypical of prokaryotes^6^. When the yeast extract concentration in the HC1 culture was increased from 0.005% to 0.05%, the growth rate (doubling times from 7 days to 6 days) did not significantly change (Fig. 1c). In contrast, observation of cells showed increases in the volume of the main cell body (from 1.2±0.01 to 1.8±0.01 µm^3^ estimated based on cell diameter increase from 1.3±0.3 to 1.5±0.3 µm) (Fig. 3d, e, and Supplementary Table 3). We further used wide-area SEM to analyze roughly 15,000 cells for changes in protrusion and vesicle production (Supplementary Fig. 5). We observed consistent increases, on a per cell basis, in volume and maximum number of protrusions (1.3 to 5.2 µm^3^ and 8 to 11, respectively), the volume and number of vesicles (0.02 to 0.11 µm^3^ and 0.4 to 3.7, respectively), and protein content (66 to 226 fg; Supplementary Table 9). Similar trends were observed for *P. syntrophicum* MK-D1 and SC1 (Extended Data Fig. 9, Supplementary Table 10, and Supplementary Note 6).

Collectively, these findings suggest that, when more carbon and energy are available, these strains allocate them primarily to the cell (that is, main cell body and protrusions) rather than to cell division. This is in direct contrast with typical prokaryotes that allocate available carbon and energy to the cell body and cell division and somewhat analogous to the biology of eukaryotes, for which size and reproduction rates are inversely related ^6,43,44^. We suspect that *Promethearchaeota*’s “aprokaryotic” focus on cell maintenance/construction over growth may have played an important role in the evolution of more complex cell structures post-FECA. Potentially related to this, for all three *Promethearchaeota* strains, only non-dividing cells produce complex actin-containing protrusions, which are a major contributor to cell size (Fig. 2 and Extended Data Fig. 3). As the number and volumes of protrusions increase when more carbon and energy are available and the protrusions are never observed in dividing cells, we suspect that, besides syntrophy, the protrusions and its contents (including actin filaments) may serve as nutrient, energy, and building block storage for later cell division.

### Strict dependency on reduced anaerobic conditions

HC1 is clearly dependent on an anaerobic lifestyle as it possesses oxygen-sensitive enzymes (*e.g.*, pyruvate- and 2-oxoacid-ferredoxin oxidoreductases); however, we also find genes encoding enzymes indicative of adaptations to a more oxidized habitat— cytochrome *bd* oxidase (Cbd)-like protein, cytochrome *c* oxidase (Cox), catalase, and superoxide dismutase. In HC1 cultures, we also verified the expression of these genes (Supplementary Table 5) and the presence of heme B (Supplementary Fig. 6), which is a cofactor for three of the above enzymes. However, based on the genome, we can predict that HC1 cannot live aerobically (Fig. 4). HC1 lacks a discernible mechanism for funneling electrons to the terminal oxidase Cox. While complex III (that is, cytochrome *bc*_1_/*b*_6_*f*) typically facilitates this, HC1 only possesses the cytochrome *b* component and lacks a corresponding cytochrome *c*, cytochrome *f*, or copper-binding electron transfer protein like plastocyanin or halocyanin for reducing Cox. While HC1 possesses proteins for funneling electrons to the quinone-dependent Cbd-like protein (*e.g.*, via a type II NADH:quinone oxidoreductase and electron transfer flavoprotein-quinone oxidoreductase), it lacks genes that would allow transfer of all reducing equivalents from peptide degradation (*i.e.*, reduced ferredoxin and NADPH) to respiration of oxygen—*e.g.*, ferredoxin:NAD^+^ oxidoreductase and transhydrogenase. Thus, oxygen reduction is, at most, a supplementary metabolism.

**Fig. 4.**
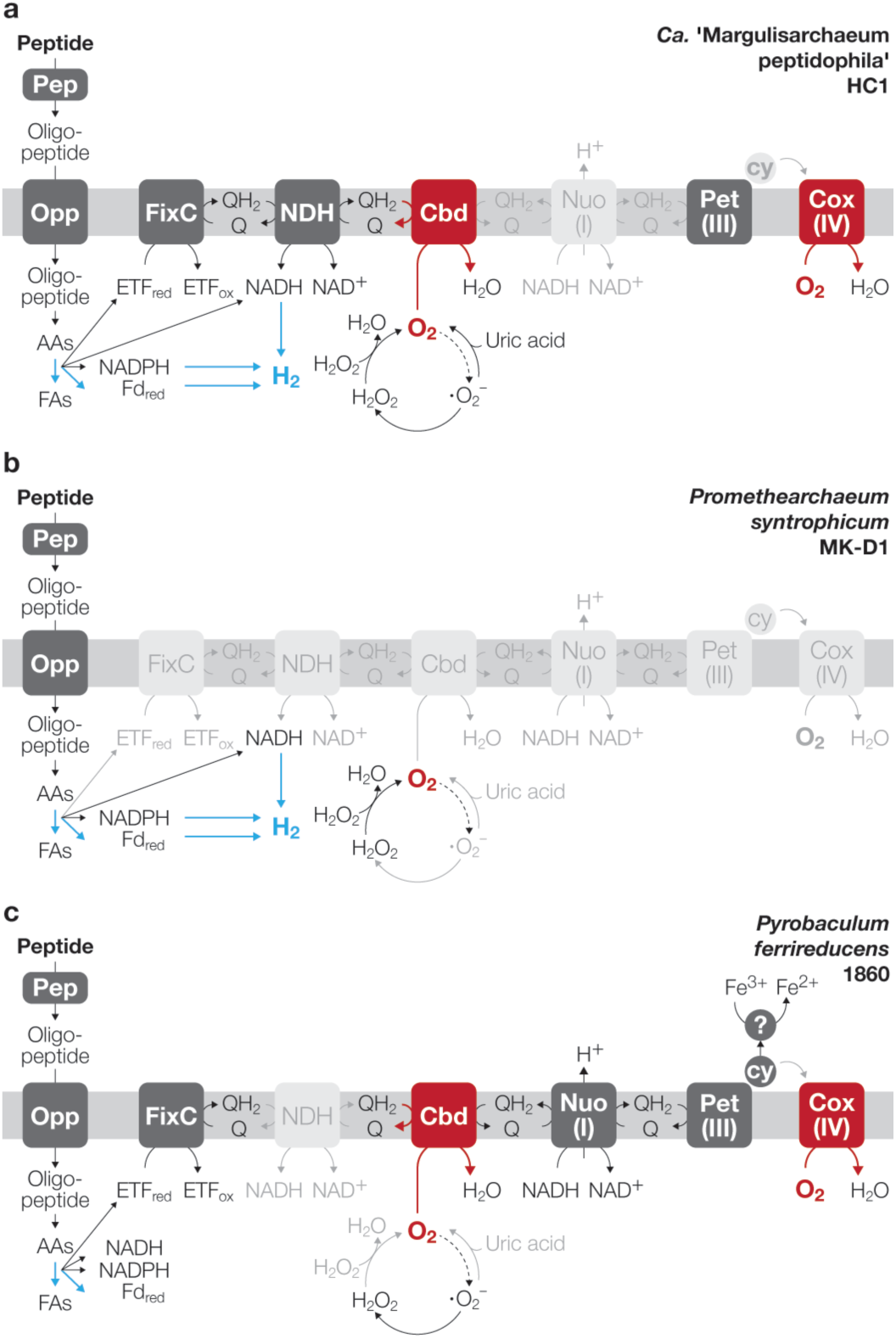
Electron transport chain and oxygen reduction-related functions of HC1, *P. syntrophicum* MK-D1, and *P. ferrireducens* 1860. **a–c,** Schematic diagrams of electron transport-chain related functions of HC1, *P. syntrophicum* MK-D1, and a representative obligately anaerobic peptidotrophic archaeon possessing terminal oxidases (*P. ferrireducens* 1860), respectively. Enzymes/reactions encoded by each strain’s genome is indicated in black, those that utilize O_2_ are indicated in red. Enzymes/reactions that each strain does not possess genes for are indicated in gray. Pathways sensitive to oxygen are indicated in blue. Abbreviations: Pep, extracellular peptidase; Opp, oligopeptide permease; AAs, amino acids; FAs, fatty acids; Fd, ferredoxin; ETF, electron transfer flavoprotein; Q, quinone; QH_2_, quinol; FixC, ETF:Q oxidoreductase; NDH, type 2 NADH:Q oxidoreductase; Nuo, type 1 NADH:Q oxidoreductase (or complex I); Cbd, cytochrome *bd* oxidase; Pet, cytochrome *bc*_1_ (or complex III); Cox, cytochrome *c* oxidase (or complex IV); cy, cytochrome *c*. For HC1, Nuo-related genes are encoded in the genome but annotated as absent because some putative subunits that would be critical for function as complex I are associated with other oxidoreductases in the genome (Supplementary Table 5).

Given that HC1 could be repeatedly sub-cultured in reducing agent-free medium with low levels of oxygen in the enrichment process, HC1 clearly has some tolerance of oxygen (Supplementary Table 1). Notably, the *Promethearchaeales* population in the same culture was no longer detectable in the first sub-culture, indicating no tolerance of oxygen. Interestingly, the purified HC1 co-cultures showed no growth in culture media containing trace levels of oxygen by (i) omitting reducing agents or (ii) supplementing the head space with gas containing 1% (v/v) oxygen. Both resulted in major decreases (roughly two orders) in HC1 cell densities (Fig. 3a). Thus, the strain is not stimulated by oxygen and can only tolerate continuous exposure to oxygen when aerobic organisms are present and presumably maintaining oxygen at trace levels (Supplementary Table 1).

In line with this, Cbd and Cox may both be used for aerotolerance rather than aerobic respiration. A previous study showed that a strict anaerobe can utilize a terminal oxidase (Cbd) to scavenge and tolerate nanomolar concentrations of oxygen, though oxygen does not stimulate growth^28^. Likewise, Cox can have affinities in the nanomolar range^45^ and our genomic survey reveals many Cox-possessing organisms described as strict anaerobes (Supplementary Table 11). This included an obligately anaerobic peptidotrophic archaeon *Pyrobaculum ferrireducens* strain 1860 possessing Cox but incapable of growing at O_2_ concentrations even as low as 0.5% (roughly 3 µM dissolved O_2_ at 90°C)^46^ (Fig. 4c). Perhaps to further support aerotolerance, HC1 possesses and expresses genes encoding enzymes for oxygen-tolerant central metabolism (pyruvate dehydrogenase) and combatting oxidative stress (*i.e.*, catalase, superoxide dismutase, thioredoxin-dependent peroxiredoxin, peroxiredoxin 2/4, and glutaredoxin-dependent peroxiredoxin; Supplementary Table 5). HC1 also possesses and expresses genes encoding proteins (*e.g.*, xanthine dehydrogenase; Xdh) for production of the antioxidant uric acid from purines, an oxidative stress mitigation strategy often associated with animals^47^. Ferritin, a key protein for reducing reactive oxygen species generation from the interaction of iron and oxygen^40^, was also among the top three highest expressed proteins in HC1’s transcriptome. Gene expressivity predicted from codon utilization^48^ also supported our interpretation that oxygen reduction is a supplementary function in the physiology of HC1 (Supplementary Table 5 and Supplementary Note 7).

As with oxygen reduction, HC1 possesses a nitrate reductase but lacks pathways for funneling all electrons from peptidotrophy to nitrate reduction. In HC1 and methanogen co-cultures supplemented with nitrate, we observed nitrate reduction activity and associated nitrite accumulation, but no stimulation of growth (Extended Data Fig. 10a). As predicted from the genome, only a small proportion of the electrons derived from peptide degradation (∼15%) were funneled into nitrate reduction and the rest were transferred to the partner methanogen. Moreover, nitrate-amended cultures showed no growth when a methanogen inhibitor was present, indicating that HC1 strictly depends on syntrophy even when performing nitrate respiration. Given that nitrate (1 mM) is inhibitory (Extended Data Fig. 10b), we suspect the nitrate reductase may serve to scavenge nitrate rather than support nitrate-respiration-driven growth. This is analogous to the terminal oxidase-supported aerotolerance discussed above (Extended Data Fig. 10c).

### Implications for eukaryogenesis

The discovery of obligately anaerobic syntrophy with potential adaptations to exposure to oxygen in ‘Hodarchaeales’ has major implications for eukaryogenesis. We here show evidence that *Promethearchaeati* species widely conserve anaerobic peptidotrophy based on analysis of gene expressivity across all archaeal species representatives (see Methods; Fig. 3f and Supplementary Table 12). Moreover, nearly all such species (89 out of 94) encoded NiFe hydrogenase gene clusters similar to either (i) those highly expressed by HC1, SC1 or *P. syntophicum* MK-D1 or (ii) the archaeal electron-bifurcating hydrogenase MvhADG-HdrABC^49^ (Fig. 3f). In detail, these NiFe hydrogenases (PF00374) form gene clusters with electron transfer subunits related to FrhG (COG1941) and at least one of NAD(P)H-flavin reductase, HdrB-like, or HdrA (COG0543, COG2048, and COG1148, respectively). The consistent coupling of organotrophy with putatively energy-conserving H_2_ metabolism indicates syntrophy is a widely conserved and ancient feature of *Promethearchaeati*, which agrees with our previous hypothesis^3^ and phylogenetic analyses tracing organotrophic H_2_ production and acetogenesis via the Wood-Ljungdahl pathway to the *Promethearchaeati* ancestor^50^. Further evidencing these organisms’ dependency on symbiosis with other organisms, *Promethearchaeota* members, including HC1 and SC1, lack many pathways for biosynthesis of amino acids, vitamins, and nucleotides (Supplementary Table 13).

In contrast, oxygen utilization via Cox is only found in ‘Heimdallarchaeia’ (Supplementary Table 12). Like HC1, metabolic reconstruction suggests that Cox-mediated oxygen reduction is a supplementary metabolism for these archaea. All Cox-possessing ‘Heimdallarchaeia’ lack complex III entirely or only possess a partial complex of unknown function (Supplementary Table 12). The absence of cytochrome *c*, cytochrome *f*, and blue copper electron transfer proteins were confirmed by a homology search using prokaryotic references with protein-level evidence and the following domains IPR009056, IPR002325, and IPR000923 respectively. Many complex III-like cytochrome *b*-possessing ‘Heimdallarchaeia’ also lack Cox. As such, we speculate that the complex III-like cytochrome *b* does not play a direct/central role in O_2_ utilization by these archaea. Similarly, some ‘Heimdallarchaeaia’ encode Cbd, but lack type 1 and 2 NADH:quinone oxidoreductases (that is, complex I and NDH-2) for electron transport. In addition, all analyzed Cox- and Cbd-possessing ‘Heimdallarchaeia’ species representatives have high-expressivity 2-oxoacid:ferredoxin oxidoreductases, a group of proteins known to be oxygen-sensitive. Thus, as with HC1, all other ‘Heimdallarchaeia’ members lack complete aerobic respiration pathways and may use terminal oxidases for aerotolerant anaerobiosis rather than oxygen respiration. However, our analyses do not reject the possibility that ‘Heimdallarchaeia’ possess uncharacterized proteins that fill in the gaps in electron transport.

To investigate whether FECA may have already had such aerotolerance, we performed phylogenetic analyses of genes encoding proteins directly related to aerotolerance and aerobic respiration (Fig. 5a-f and Supplementary Figs. 7–11). For Cox subunits 1 and 3 (COX1 and COX3) and heme O synthase (COX10) (Fig. 5b-d), ‘Heimdallarchaeia’ form monophyletic groups with ‘Hodarchaeales’ sequences in basal positions and ‘Heimdallarchaeales’ nested within. This suggests first acquisition by ‘Hodarchaeales’ and subsequent horizontal gene transfer to the latter. A similar relationship was observed for Xdh (Fig. 5e), which may support aerotolerance via production of uric acid as a reactive oxygen species scavenger^47^. Thus, the ancestor of ‘Hodarchaeales’ may have possessed some aerotolerance. If FECA emerged as a sister clade to ‘Hodarchaeales’, FECA may have already possessed such aerotolerance (Fig. 5g). If FECA diverged earlier as a sister clade to ‘Heimdallarchaeia’, FECA would have lacked aerotolerance mediated by the above genes (Fig. 5g). Collectively, our data indicates that FECA was an archaeon equipped with anaerobic syntrophic peptidotrophy and, potentially, aerotolerance. While one ‘Heimdallarchaeia’ clade nearly fully conserves Cox (’Kariarchaeaceae’) and are likely better adapted to oxygen than other *Promethearchaeota*, relevance of their phenotype to FECA remains unclear given that they horizontally acquired heme O synthesis and Cox from ‘Hodarchaeales’ presumably after FECA diverged (Fig. 5b–d, g).

**Fig. 5.**
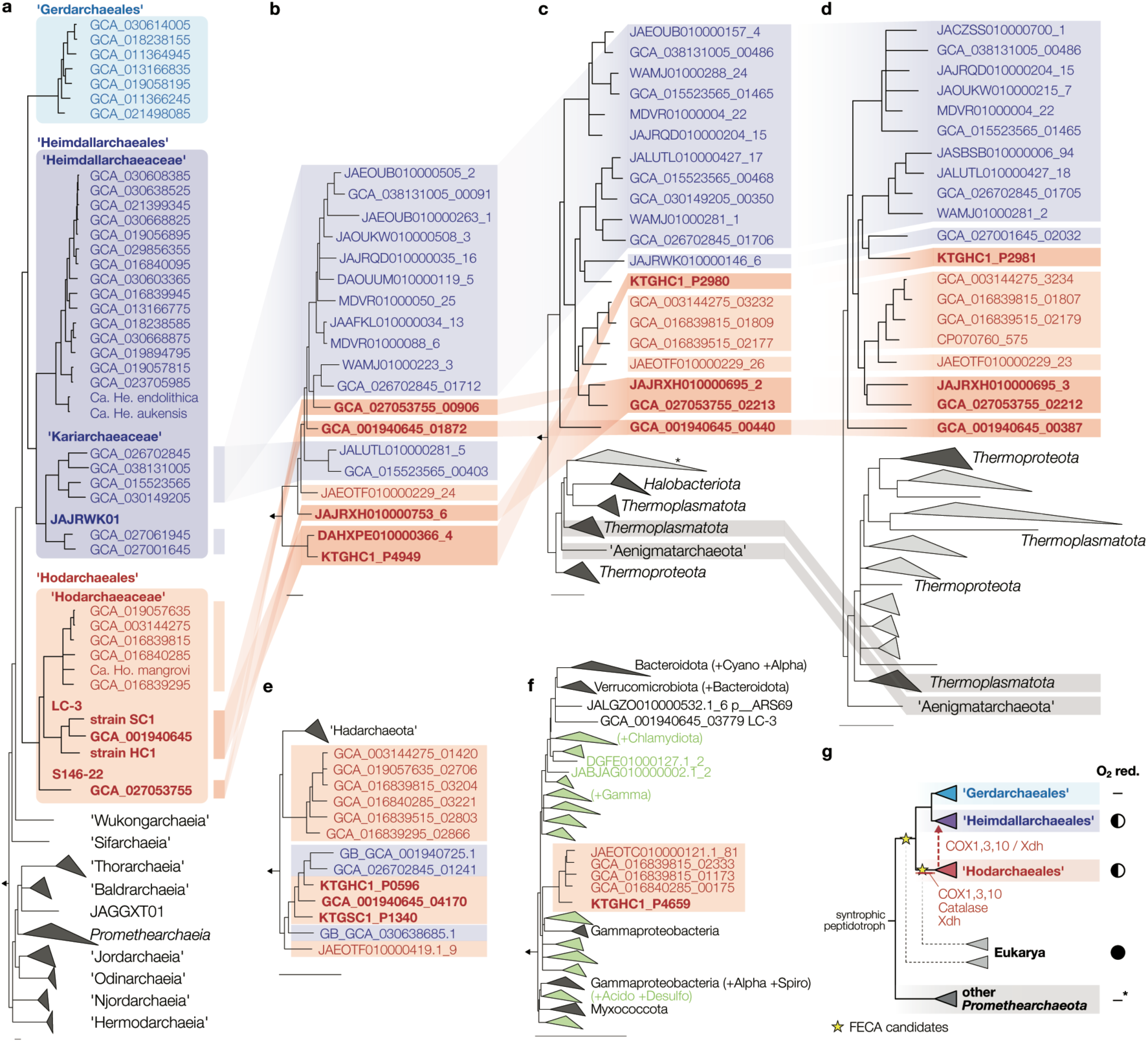
Phylogeny of HC1 and SC1 and genes related to oxygen metabolism. **a,** Maximum-likelihood estimation of the phylogeny of *Promethearchaeati* based on a concatenated alignment of 47 marker genes using IQTREE2. **b–f**, Maximum-likelihood estimations of the phylogeny of (**b**) heme O synthase (COX10), (**c**) cytochrome *c* oxidase subunit 1 (COX1), (**d**) subunit 3 (COX3), (**e**) xanthine dehydrogenase (Xdh), and (**f**) catalase. For each tree calculation, optimal models from C10, C20, C30, C40, C50, and C60 (+LG+G+F) were chosen using ModelFinder. Branch support was calculated using ultrafast bootstrap and BOOSTER. All branches with support less than 0.95 are collapsed. Sequences belonging to ‘Heimdallarchaeales’, ‘Gerdarchaeales’, and ‘Hodarchaeales’ are colored dark blue, light blue, and red respectively. Clades with alphaproteobacterial sequences in basal positions are indicated in green. Full phylogenetic trees are shown in Supplementary Figures 7–11. **g**, A schematic diagram showing the acquisition and transfer of aerobiosis-related genes in *Promethearchaeati* and their relationship with points in the tree previous studies propose eukaryotes may have diverged from *Promethearchaeati* (that is, FECA). The presence of members with the genomically predicted capacity to reduce oxygen but not live aerobically (missing discernible electron transfer pathways connecting organotrophy to oxygen reduction [O_2_ red.]) is indicated with a half-filled circle. Absence of oxygen reduction in a lineage is indicated with a hyphen (*some *Promethearchaeati* species outside of ‘Heimdallarchaeia’ possess partial terminal oxidases of unknown function).

Phylogenetic analysis of catalase showed that ‘Hodarchaeales’ acquired the protein horizontally from *Alphaproteobacteria* (Fig. 5f). This is the first evidence for horizontal gene transfer between these two respective lineages presumably closely related to FECA and protomitochondrion. Whether this has a direct relationship with eukaryogenesis remains unknown. However, this suggests that the two lineages co-existed in an anoxic habitat exposed to oxygen where acquisition of catalase would have been advantageous for ancient ‘Hodarchaeales’. This aligns well with (i) behavior observed in our reducing agent-free enrichment cultures containing HC1 (Supplementary Table 1), (ii) the potential aerotolerance of ancestral ‘Hodarchaeales’, and (iii) a model for eukaryogenesis we previously proposed (the “E^3^ model”)^3^. In addition, the aerotolerance of HC1 exclusively observed in cultures containing oxygen-respiring organisms is analogous to oxygen-centric symbiosis between FECA and the aerobic protomitochondrion theorized to be important to the rise of eukaryotes.

## Conclusion

In this study, we isolated two strains from the archaeal lineage most closely related to FECA—’Hodarchaeales’—and elucidated their biology. The consistency in physiology and cell structure between *P. syntrophicum* MK-D1^2,3^, ‘L. ossiferum’^4^, and strains HC1 and SC1 suggests the above features are ancestral to and conserved across *Promethearchaeota*, which is further supported by genomic analyses. Discovery of a clear connection between syntrophy and cell structure (that is, the protrusions) further supports the ancestral coincidence of these features. Thus, we can deduce that FECA was an anaerobic, syntrophic, peptide-degrading, mesophilic archaeon that had simple cell structure with branching protrusions and no endomembrane system. Phylogenetic analyses suggest FECA may have also had low aerotolerance but did not respire oxygen, like HC1 (Fig. 5g). This strongly indicates that the prokaryote-eukaryote transition was steep in both cell structure and physiology—from anaerobic to aerobic and from simple to complex internal cell structure. Slow growth is also conserved across all cultured strains and we suspect this may have played an important role in eukaryogenesis, including fixation of the bacterial endosymbiont, replacement of archaeal with bacterial lipids, and “bacterialization” of the genome (Supplementary Note 8).

While the extremely small cell size and slow growth rate of *P. syntrophicum* MK-D1, the only publicly available *Promethearchaeati* strain^2,3^, had posed a major obstacle, efforts in cultivation here led to the isolation of two strains that have larger cell sizes, higher growth rates, shorter lag phases, and a closer relationship to eukaryotes. We believe these make the strains more suitable as model organisms for exploring the biological properties of *Promethearchaeati* and can catalyze further breakthroughs in uncovering the transition from simple to complex life.

## Methods

No statistical methods were used to predetermine sample size. The experiments were not randomized. The investigators were not blinded to allocation during experiments and outcome assessment.

### Sample collection and culturing

The biofilm samples were collected from a gas production well, KTG3, in the Southern Kanto gas field (Chiba Prefecture, Japan). The biofilm was located on a wall of the separator discharge outlet at a depth of 20–40 cm below the water surface. Samples were taken after eruption of brine on August 29th, 2019, July 16th, 2020 and May 11th, 2023 (Extended Data Fig. 1). The gas field is located in an old marine forearc basin and is based on subsurface gas reservoirs in turbidite sandstones, making it geologically and geochemically relevant to past seawater and marine sediment^51^. The brine from the well KTG3 is rich in organic matter, iodine, and iron. Detailed site information has been described previously^33^ and geochemical characteristics of the brine in the well KTG3 are provided in Extended Data Fig. 1b. The collected biofilm sample was placed in a 100-ml glass bottle filled with brine (no head space) and brought back to our laboratory under refrigerated conditions. Samples were also inoculated in 50-ml glass vials containing anaerobic medium onsite (Supplementary Note 2). Our iTag analysis showed that the biofilm contained abundant and diverse microorganisms, mostly affiliated with uncultured or cultivation-limited microbial groups, including *Promethearchaeota*. To isolate new *Promethearchaeota* strains from the samples, we employed a strategic batch-type cultivation^31^ that incorporated previous knowledge and our expectations for this archaeal group (Supplementary Note 1). Detailed information about the cultivation, including the medium composition and purity checks for the HC1 and SC1 cultures, is described in the Supplementary Methods. Unless otherwise indicated, the HC1 and SC1 cultures were grown in casamino acids–peptone–yeast extract medium (each at 0.005%, w/v) and casamino acids–powdered milk–polypeptone peptone–yeast extract medium (each at 0.005%, 0.005%, 0.01% and 0.05%, w/v), respectively, with inoculum size of 16% v/v for HC1 and 20% v/v for SC1.

### Growth monitoring using qPCR and iTag analysis

DNA extraction, PCR mixture preparation, and qPCR analysis were performed as described previously^3,52^. PCR primers for qPCR used in this study were designed using the probe design tool of ARB program^53^. The specificity of new primers was confirmed using BLAST and the ARB-SILVA database (https://www.arb-silva.de/)^54^. The primers and PCR conditions are shown in Supplementary Table 14. The doubling times of cultured archaea were calculated based on the semilogarithmic plot of the qPCR data. For iTag analysis, a universal primer pair 530F/907R, which contained overhang adapters at 5’ ends, was used^55^. The procedures used for library construction, sequencing, and data analysis were described previously^3^. All samples for data acquisition were checked for growth and microbial composition by qPCR and iTag before or after the experiments.

### Evaluation of growth temperature and substrate usage

To evaluate the temperature range for growth, a pure co-culture of HC1 and a tri-culture of SC1 were incubated at the range of 10 to 45 or 50°C. All incubations were performed in triplicate. After a certain period of incubation, 16S rRNA gene copy numbers of each strain were evaluated by qPCR.

To examine substrate utilization of the cultured archaeal strains, the basal media were supplemented with each substrate sterilized either by autoclaving or filtration. We performed two successive transfers under the same cultivation conditions and conducted qPCR targeting the archaeal strains’ 16S rRNA genes in both the first and second subcultures. All incubations were performed in duplicates at the optimum growth temperature for each strain.

To assess the effect of non-proteinaceous substrates on growth of archaeal strains, we added individual substrates to culture media containing casamino acids–peptone–yeast extract for HC1 and casamino acids–powdered milk–polypeptone peptone–yeast extract for SC1. Each cultivation condition was performed in duplicate.

### Epifluorescence microscopy

An epifluorescence microscope (BX53, Olympus) equipped with a phase-contrast objective lens and a color CCD camera system (DP80 or DP75, Olympus) was used for routine microscopic observation and video recording of the microbial cells. When it was necessary to obtain images without cell movement due to Brownian motion, agarose-coated glass slides were used^56^.

### SEM

For SEM analysis, archaeal cells were fixed by adding 2.5% (w/v) glutaraldehyde and incubating for 2 h under anoxic conditions at their optimum growth temperatures and then overnight at 4°C. The sample preparation procedure was performed as described previously^2^. The samples were observed under an extreme high-resolution focused ion beam (FIB)-SEM (Helios G4 UX, ThermoFisher Scientific) operated at 1kV.

To evaluate the volumes and numbers of HC1 cell bodies, protrusions, and membrane vesicles under different yeast extract concentrations, we acquired wide-area SEM images using Maps 2.5 (Thermo Fisher Scientific), which was installed in the FIB-SEM operation system. Wide-area images were obtained by stitching 100 images into a larger 10 × 10 image, with each image taken with an acceleration voltage of 1 kV, a beam current of 13 pA, a field of view of 36.3 µm × 24.2 µm, and an image resolution of 23.6 nm/pixel. The acquired wide-area SEM images were then analyzed using AMIRA 3D Pro 2024.1 and 2024.2 software (Thermo Fisher Scientific). First, the images were denoised by applying curvature-driven diffusion^57^. Next, HC1 cell bodies, protrusions, membrane vesicles, and *Methanocalculus* cells were segmented with the aid of a deep learning module based on PyTorch^58^ and Keras (https://github.com/fchollet/keras). The deep learning model created from 2000 × 2000 pixels was adapted to the full images, followed by manual modification and watershed segmentation. Each segmented component, except for protrusions, was labeled and analyzed the area and perimeter. Protrusions were skeletonized using the Auto Skeleton module^59^, and the number of fiber segments and their curved length were measured after excluding looped fibers or those shorter than 10 pixels.

### TEM

In the preparation of ultrathin sections of archaeal cells for TEM, we followed a previously established protocol^2^. The section samples were observed using a conventional TEM (Tecnai 20, FEI) at an acceleration voltage of 120 kV. Detailed information about observations of HC1’s membrane vesicles and thin filaments on the surface of SC1’ protrusions is described in the Supplementary Methods.

### QFDE-EM

For QFDE-EM observations, archaeal cultures (440 ml of HC1 culture, and 25 ml of SC1 culture) were concentrated to approximately 5 ml through filtration using a 0.22-µm-pore-size polyethersulfone (PES) filter unit (Corning) and transferred to a glass vial in an anaerobic chamber (95:5 [v/v] N_2_:H_2_ atmosphere; COY Laboratory Products). To remove the H_2_, which is inhibitory to the strains, this vial’s headspace was replaced with N_2_/CO_2_ gas (80:20, v/v) via flushing. The subsequent procedure was performed as described previously^2,60^.

### Cryo-EM

Prior to the electron microscopic observation, 2 ml of culture liquid was taken from a culture bottle using a syringe, and the culture liquid was concentrated to approximately 50 µl by centrifugation at 20,400×g for 10 min at 20°C. Then, 3 µl of the concentrated liquid culture was applied onto a glow-discharged Quantifoil Mo grid R2/1 (Quantifoil MicroTools) after mixing with 15 nm gold colloids (Electron Microscopy Sciences) and plunged-frozen in liquid ethane using a Vitrobot Mark IV (Thermo Fisher Scientific) at 4°C and 95% humidity. The frozen grids were screened on a 626 side-entry cryoholder (Gatan) with a 200kV JEM2200FS electron microscope (JEOL). Suitable grids were subjected to cryo-electron tomography in a 300 kV Titan Krios G4 electron microscope (Thermo Fisher Scientific) equipped with a bottom-mounted Selectris X energy filter (slit width: 5 eV) and a Falcon 4i direct detector camera. Tile series of each specimen were collected at a nominal magnification of ×19,500, resulting in an image resolution of 4.633 Å per pixel, and a dose symmetric tile scheme was performed between ± 60° at 2° using Tomography software implemented on a TITAN Krios G4, resulting in a total electron dose of 91.5 electrons per Å^2^. The tilt series was aligned with gold fiducials, and tomograms were reconstructed using filtered back projection or SIRT in the IMOD software^61^ with an image binning of 5. Segmentations of HC1’s cellular components were performed using AMIRA 3D Pro software (Thermo Fisher Scientific). Cellular membranes and granular structures were segmented manually with the help of a deep learning module, and actin filaments were segmented manually.

### Immunofluorescence staining using lokiactin-specific antibodies

The protocol for antibody preparation and observation by immunofluorescence for lokiactin generally followed that of Rodrigues-Oliveira et al^4^. The lokiactin antibodies were made using peptides specific for the lokiactin sequences of HC1 and SC1 (Supplementary Fig. 2). The regions of the peptides are the same region as the peptides for *Ca*. ‘L. ossiferum’^4^. We made three antibodies in total (that is, ab1 for HC1, ab1 for SC1, and ab2 for both strains), and all of them were created using the service of Cosmo Bio Co., Ltd. and validated by ELISA assays. Detailed procedures of western blotting and immunofluorescence staining are described in the Supplementary Methods.

### Hybridization chain reaction (HCR)-FISH

Cells were fixed under anoxic conditions by adding paraformaldehyde solution (methanol-free, formaldehyde solution, 16% w/v, Electron Microscopy Sciences) to a final concentration of 4% (v/v) directly into the culture bottles and incubating for 2–3 hours at 4°C. After fixation, cells were washed with phosphate-buffered saline (PBS; 153.8 mM NaCl, 2.7 mM Na_2_HPO_4_, 1.5 mM KH_2_PO_4_ [pH 7.2]). Subsequently, cells were stored in 50% ethanol with PBS at -30°C until use in subsequent experiments. The design of specific probes and HCR-FISH were performed in accordance with previously described protocols^3,62^. The 16S rRNA-targeted oligonucleotide probes used in this study are listed in Supplementary Table 15. Cells were counterstained with DAPI in the ProLong Diamond Antifade Mountant with DAPI (Thermo Fisher Scientific). FISH images were acquired using a STELLARIS STED confocal microscope (Leica) with a ×100/1.4 NA oil-immersion or ×93/1.3 NA glycerol-immersion objective lens at 2 or 6 × zoom, line scanning frequency of 400 or 700 Hz, and line average of 3. Fluorescence signals were detected in confocal mode at 405, 499, and 553 nm for excitation, and some images were deconvolved using the adaptive Lightning strategy in LasX software (Leica) when necessary. The 3D movie was constructed from 27 sections of 8.17 μm thick using the LasX software. Line intensity profiles were obtained using LasX software for each of the DAPI and Alexa Fluor 488 channels along a line through the center of the target cell.

### CLEM

Correlative light-electron microscopy was conducted as previously described^63^ with minor modifications. The detailed procedure is described in the Supplementary Methods.

### Chemical analyses

To measure peptides and proteins, culture liquid samples were filtered through a 0.2 µm pore-size polytetrafluoroethylene (PTFE) filter unit (Merck Millipore) before sample preservation and analysis. The molecular weight distribution of peptides and proteins was determined using an ultra-performance liquid chromatography equipped with ACQUITY UPLC Protein BEH SEV column (Waters) as described previously^64^.

Methane concentrations in the headspace of culture bottles were measured by gas chromatography as described previously^3^.

Measurement of heme B was carried out according to the previously reported method^65^. For this measurement, archaeal cultures (440 ml of HC1, and 900 ml of *P. syntrophicum* MK-D1) were first concentrated to about 5 ml using a 0.22-µm-pore-size polyethersulfonate (PES) filter unit (Corning) on a clean bench. The concentrated cell suspensions were then further concentrated by pelleting the cells through centrifugation at 20,400×g for 10 min at 4°C. The harvested cells were washed once with their respective basal media without energy substrates.

Nitrate and nitrite concentrations were measured by high-performance liquid chromatography (HPLC) using a TSKgel SAX column (Tosoh Co., 35 mm × 4.6 mm i.d.) with 3% (w/v) NaCl; as the eluent. The column temperature was maintained at 40 °C, and detection was performed with a UV-VIS detector (GL-7451, GL Science), as described previously^66^.

Iron concentrations in the brine were determined by quadrupole inductively coupled plasma mass spectrometry (ICP-MS; iCAP Qc, Thermo Fisher Scientific). To prevent metal precipitation before measuring total iron concentrations, brine samples were acidified with high-purity concentrated HNO_3_ (TAMAPURE AA-100, Tama Chemical Co. Ltd.) at a final concentration of 1.5 M. For ICP-MS analysis, the water sample was diluted with 0.3 M HNO_3_, and four internal standards (Be, Sc, Y, and In) were added to correct instrument drift. Mass interferences were eliminated using the kinetic energy discrimination mode with H_2_ reaction gas for high-sensitivity ^56^Fe analysis.

### Genomic, transcriptomic and proteomics analyses

Genomic DNA from HC1 and SC1 was extracted according to a previous protocol^52^. For whole genome short-read sequencing, paired-end libraries were prepared with KAPA Hyper Prep Kit (Roche) according to the manufacturer’s instructions. To supplement the genome assembly, an Illumina mate-pair library for HC1 and a Nanopore library for SC1 were constructed and sequenced using an Illumina MiSeq and an Oxford Nanopore PromethION platform, respectively. The mate-paired library with an average insert size of 3 kb was prepared with Nextera Mate Pair Library Preparation kit (Illumina). Raw short reads were trimmed using Trimmomatic ver. 0.39^67^ to remove adapters and low-quality bases (Phred score < Q20) with a minimum read length of 100 bp. Linker sequences in mate-paired reads were removed using NextClip v1.3.1^68^. Short-read assemblies were generated with CLC Genomic Workbench v23 (Qiagen) and SPAdes v3.15.3^69^. The resulting assemblies were manually merged and optimized to produce longer contigs. Scaffolding was performed with BESST v2.2.8^70^ for mate-pair and LRScaf v1.1.8^71^ for long reads. Gaps within the scaffolds were filled with short reads using GapFiller v1.10^72^. Coding sequences (CDSs) were predicted with Prodigal v2.6.3^73^. Deduced amino acid sequences were subjected to BLASTP search against the NCBI non-redundant protein database and the KEGG GENES database, and functional annotations were manually assigned based on the KEGG Orthology database. Prediction of rRNA and non-coding RNA genes was performed with Rfam v13.0^74^, and tRNA genes were identified using tRNAscan-SE v1.3.1^75^.

For RNA-based sequencing analysis, total RNA was extracted from HC1 and SC1 cells harvested during their exponential growth phase. Specifically, cells were collected from a 25-ml HC1 culture grown for 25 days and a 50-ml SC1 culture grown for 51 days. To concentrate the cells, the cultures were first filtered through a 0.22-µm-pore-size PES filter unit (Corning) on a clean bench, reducing the volume to approximately 4 ml, and then centrifuged at 20,400×g for 10 min at 4°C for further concentration. RNA was extracted using the RNeasy Mini Kit (Qiagen) according to the manufacture’s instructions and subsequently concentrated with the RNA Clean & Concentrator-5 Kit (Zymo). RNA quality and quantity were assessed using an Agilent 2100 Bioanalyzer system with an RNA Pico kit (Agilent Technologies). Library preparation was performed using TruSeq Stranded mRNA Library Prep Kit and TruSeq RNA CD Index Plate (Illumina). The constructed cDNA library was sequenced on an Illumina HiSeqX platform. Raw RNA-seq reads were processed using Trimmomatic ver. 0.39^67^ to trim low-quality and adapter sequences. The trimmed reads were aligned to the reference genome using Bowtie2 v2.3.5.1^76^, and gene expression levels were quantified with featureCounts v1.6.385^77^ using the corresponding gene annotations. Gene expression counts were normalized using the Python package RNAnorm v2.1.0 (https://github.com/genialis/RNAnorm).

For proteomic analysis, HC1 was cultured for 30 days in a casamino acids-peptone-yeast extract medium at two different yeast extract concentrations (0.005% and 0.05% [w/v]), using the same parent culture as the inoculum. SC1 was cultured for 50 days in a casamino acids-powdered milk-polypeptone peptone-yeast extract medium. *P. syntrophicum* MK-D1 was cultured for 105 days in a casamino acids–powdered milk medium. After confirming growth by qPCR, cells were concentrated using a PES filter unit followed by centrifugation, as done for the samples used in western blotting analysis. The collected cell samples were washed once with the basal media lacking energy substrates, then suspended in 200 µl of 100 mM triethylammonium bicarbonate (pH 8.6) containing 2 mM phenylmethylsulfonyl fluoride. Cells were disrupted by sonication at 2°C using a Q700 sonicator (QSonica) operating at 40 kHz. The protein concentration of the cell-free extract was determined by a Qubit fluorometer (Thermo Fisher Scientific). All subsequent procedures, including liquid chromatography-tandem mass spectrometry (LC-MS/MS) data acquisitions and data analysis, were performed as described previously^78,79^.

### Glycoproteome analysis

For glycoproteome analysis, the same culture samples as those used for proteome analysis were employed. The procedure was identical up to the point of cell disruption by sonication, after which half of the protein extract samples was allocated for glycoproteome analysis. Subsequent sample preparation, LC-MS/MS analysis, and data processing were performed as previously described^42,80^.

### Phylogenetic analyses

*Promethearchaeota* genomes were collected from the Genome Taxonomy Database (GTDB) release 220^81^ and GenBank. Genomes not included in GTDB r220 were classified into species-level lineages defined in GTDB or new species based on sequence distances estimated using MASH v2.3^82^. Coding sequences were predicted using Prokka 1.14.6^83^ with default parameters and annotated with eggnog-mapper v2.1.12 and the eggnog v5.0 database using default parameters^84^. For constructing a genome-based phylogeny of *Promethearchaeota*, the following single copy marker genes defined by GTDB were used for phylogenetic tree construction (Supplementary Table 16). Maximum likelihood estimation of the phylogeny was performed using IQTREE v2.3.5 with ultrafast bootstrapping^85^, a concatenated alignment of these marker genes trimmed using BMGE v1.12 (-m BLOSUM30 -g 0.67 -b 3)^86^, and a universal distribution model (UDM0064 LCLR) as previously described^87^. Maximum likelihood estimation of the phylogeny for the 16S rRNA gene was performed using a sequence alignment generated with MAFFT (--maxiterate 1000 --localpair)^88^, the optimum model selected by ModelFinder (SYM+R6)^89^, and regular bootstrapping. Bootstrap values were recalculated using BOOSTER v0.1.2^90^.

For maximum likelihood estimation of the phylogeny of target functional genes, we first collected relevant sequences by using amino acid sequences obtained from *Promethearchaeota* as the query and searching against genes belonging to the same protein family collected from GTDB r220 species representatives^81^. For this, the MMseqs2 v16.747c6 search command^91^ was used with 3 iterative searches with different sensitivities from sensitivity level 4 to 8.5. A sequence identity cutoff of 0.3 and coverage cutoff of 0.7 was used. To select representative sequences for tree construction, we (i) clustered the collected sequences using the MMseqs2 cluster command^91^ with a sequence identity cutoff of 0.6 and coverage cutoff of 0.7, (ii) constructed a rough tree using FastTreeMP v2.1.11^92^ with an alignment generated with MAFFT v7.525^88^ and trimmed with trimAl v1.5.0 (-gt 0.1)^93^, and (iii) subsampled phylogenetically representative sequences from the tree using Parnas v0.1.6^94^. We performed protein structure-informed alignment of the representative sequences using MAFFT-DASH (--localpair --maxiterate 100 --dash)^95^ and trimmed the alignment using trimAl (-gt 0.3). Finally, we used this as an input for maximum likelihood estimation of the phylogeny using IQTREE v2.3.5 with ultrafast bootstrapping^85^ and ModelFinder^89^ to identify the best-fitting model among C10, C20, C30, C40, C50, and C60 (LG+CXX+G+F). Bootstrap values were recalculated using BOOSTER v0.1.2^90^.

### Metabolic reconstruction and codon analysis

Gene expressivity was calculated using cordon^48^ using ribosomal proteins as a reference and default settings. Peptidotrophy was annotated based on the presence of extracellular peptidases, glutamate dehydrogenases, and glycine dehydrogenases with high expressivity (within top 25% of each genome), all consistently found in isolates specialized in peptide degradation (strains MK-D1, HC1, SC1, *Coprothermobacter proteolyticus*, *Necrotrophus endoclepta*, *Porphyromonas gingivalis*, and *Salinivirga cyanobacteriivorans*). Anaerobic peptidotrophy was annotated if high-expressivity 2-oxoacid:ferredoxin oxidoreductases and FeFe or NiFe hydrogenases were also present. Those with NiFe hydrogenase gene clusters like those highly expressed by syntrophic *Promethearchaeota* isolates or archaeal electron-bifurcating hydrogenase MvhADG-HdrABC were further annotated as syntrophic. Aerobic peptidotrophy was annotated based on the presence of a high-expressivity Cox (within top 50% of each genome) along with a complete complex III (cytochrome *b*/*b*_6_ with a cytochrome *c*, Rieske FeS protein, and/or copper-binding electron carrier like plastocyanin or halocyanin). Aerotolerant peptidotrophy was annotated based on the presence of a high-expressivity Cbd with type 1 or 2 NADH:quinone oxidoreductase but lacking the ability to transfer all electrons from peptide degradation to O_2_ reduction. Different expressivity thresholds were used for peptide metabolism and terminal oxidases based on the observation that expressivity tends to be in the 25-50% range for archaea known to be aerobic (that is, *Halobacteriales*).

### Data availability

Genomes for *Ca*. M. peptidophila HC1, *Ca*. F. multiprotrusionis SC1, and *Methanocalculus* sp. strains MC2 and MC3 are available under BioSample accession numbers SAMD00653744, SAMD00736000, SAMD00653745, and SAMD00736001, respectively. The iTag sequence data was deposited in BioProjects PRJDB19594, PRJDB19596 and PRJDB19597 with SRA accession numbers DRR620266–DRR620297. The 16S rRNA gene sequences of ‘Hodarchaeales’ HC2 and ‘Heimdallarchaeaceae’ HC4 obtained from enrichment cultures were deposited in the DDBJ/EMBL/GenBank database under accession numbers LC859055 and LC859056, respectively. The RNA-Seq data of HC1 and SC1 were deposited in BioProject PRJDB19599 with SRA accession number DRR620298 and BioProject PRJDB19601 with SRA accession number DRR620299, respectively. The cryo-electron tomograms have been deposited in the EMDB with accession codes EMD-63316 and EMD-63317 for HC1 and EMD-63318– EMD-63320 and EMD-63267 for SC1. The proteome and glycoproteome data, including all LC-MS/MS raw files (.raw) and complete protein identification lists (.xlsx), have been deposited in the ProteomeXchange Consortium via jPOSTrepo^96^ under the data set identifiers PXD059994 (JPST003561) for HC1, PXD059995 (JPST003567) for SC1, PXD051006 (JPST003012) for *P. syntophicum* MK-D1.

## Acknowledgements

We thank Kanto Natural Gas Development Co. Ltd. for providing the biofilm samples, especially D. Murai, K. Kawano, N. Yokota, and M. Uzawa for their assistance in sampling the gas production well. We also grateful to A. Agata and K. Komiyama for assistance with HCR-FISH analysis; Y. Zhang for help with immunofluorescence staining experiment; M. Isozaki for help with cultivation experiments; Y. Yoshida-Takashima for help with membrane vesicle sample preparation; E. Ito for support in constructing 3D segmentation images; C. Song, Y. Kayama, R. N. Burton-Smith, and M. Ikeda for assistance with cryo-EM observations; R. C. Robinson, Y. Senju, H. Tamaki, S. E. McGlynn, and W. F. Martin for valuable advice and discussions. This study was partially supported by Japan Society for the Promotion of Science (JSPS) Grant-in-Aid for Scientific Research 19H01005 and 22H04985 to H.I., M.K.N., S.I., Y.M., and Y.Takano and 24H00582 to H.I and M.K.N, the Moore–Simon Project on the Origin of the Eukaryotic Cell (GBMF9743) to H.I. and M.K.N, and research grant L-2024-1-003 from the Institute for Fermentation, Osaka to M.K.N. This work was also supported by JSPS KAKENHI Grant Number JP22H04926, Grant-in-Aid for Scientific Research on Innovative Areas – Platforms for Advanced Technologies and Research Resources “Advanced Bioimaging Support” and the Cooperative Study Program of National Institute for Physiological Sciences (23NIPS201 and 24NIPS201). Additionally, this study was partially supported by the Research Support Project for Life Science and Drug Discovery (Basis for Supporting Innovative Drug Discovery and Life Science Research [BINDS]) from AMED under Grant Number JP24ama121005.

## Author contributions

H.I. and M.K.N conceived and led the study. H.I., S.I., and Y.S. carried out the biofilm sampling. H.I., S.I., M.Miyazaki, M.O., Y.S., and S.Sakai conducted cultivation and culture-based experiments. M.K.N. performed metabolic reconstruction, phylogenetic analyses and codon analysis. M.K.N. and Y.Takaki performed genome analysis. H.I., T.I., Y.M., K.M., M.Miyata, M.O., S.O., Y.S. Y.O.T., and K.U. undertook the microscopy and imaging work. M.O. and Y.S. performed qPCR, 16S rRNA gene analysis and DNA/RNA sequencing. Y.H. Y.I. M.Miyazaki. Y.Takano, E.T., and T.Y. performed chemical analyses. S.N. and S.Shimamura conducted proteome and glycoproteome analyses. H.I., M.K.N., S.I., K.M., S.N., S.O., S.Shimamura, Y.Takaki and K.T. conducted data interpretation. H.I. and M.K.N. wrote the manuscript with input from all co-authors. All authors have read and approved the manuscript submission.

## Author Information

The authors declare no competing financial interests. Correspondence and requests for materials should be addressed to H.I. or M.K.N.

## Additional Information

Supplementary Tables, Supplementary Videos, and Supplementary Video legends are available at figshare (https://doi.org/10.6084/m9.figshare.28480589).

## References

1. Vosseberg, J. et al. The emerging view on the origin and early evolution of eukaryotic cells. Nature 633, 295–305 (2024).

2. Imachi, H., et al. *Promethearchaeum syntrophicum* gen. nov., sp. nov., an anaerobic, obligately syntrophic archaeon, the first isolate of the lineage ‘Asgard’ archaea, and proposal of the new archaeal phylum *Promethearchaeota* phyl. nov. and kingdom *Promethearchaeati* regn. nov. Int. J. Syst. Evol. Microbiol. 74, 006435 (2024).

3. Imachi, H. et al. Isolation of an archaeon at the prokaryote-eukaryote interface. Nature 577, 519–525 (2020).

4. Rodrigues-Oliveira, T. et al. Actin cytoskeleton and complex cell architecture in an Asgard archaeon. Nature 613, 332–339 (2023).

5. Eme, L. et al. Inference and reconstruction of the heimdallarchaeial ancestry of eukaryotes. Nature 618, 992–999 (2023).

6. Kempes, C. P., Dutkiewicz, S. & Follows, M. J. Growth, metabolic partitioning, and the size of microorganisms. Proc. Natl. Acad. Sci. USA 109, 495–500 (2012).

7. Martin, W. F., Garg, S. & Zimorski, V. Endosymbiotic theories for eukaryote origin. Philos. Trans. R. Soc. B: Biol. Sci. 370, 20140330 (2015).

8. Eme, L., Spang, A., Lombard, J., Stairs, C. W. & Ettema, T. J. G. Archaea and the origin of eukaryotes. Nat. Rev. Microbiol. 15, 711–723 (2017).

9. López-García, P. & Moreira, D. The symbiotic origin of the eukaryotic cell. C. R. Biol. 346, 55–73 (2023).

10. Donoghue, P. C. J. et al. Defining eukaryotes to dissect eukaryogenesis. Curr. Biol. 33, R919–R929 (2023).

11. Spang, A. et al. Complex archaea that bridge the gap between prokaryotes and eukaryotes. Nature 521, 173–179 (2015).

12. Zaremba-Niedzwiedzka, K. et al. Asgard archaea illuminate the origin of eukaryotic cellular complexity. Nature 541, 353–358 (2017).

13. Williams, T. A., Cox, C. J., Foster, P. G., Szöllősi, G. J. & Embley, T. M. Phylogenomics provides robust support for a two-domains tree of life. *Nat*. Ecol. Evol. 4, 138–147 (2019).

14. Liu, Y. et al. Expanded diversity of Asgard archaea and their relationships with eukaryotes. Nature 593, 553–557 (2021).

15. Akıl, C. & Robinson, R. C. Genomes of Asgard archaea encode profilins that regulate actin. Nature 562, 439–443 (2018).

16. Survery, S. et al. Heimdallarchaea encodes profilin with eukaryotic-like actin regulation and polyproline binding. *Commun*. Biol. 4, 1024 (2021).

17. Hatano, T. et al. Asgard archaea shed light on the evolutionary origins of the eukaryotic ubiquitin-ESCRT machinery. Nat. Commun. 13, 3398 (2022).

18. Akıl, C. et al. Insights into the evolution of regulated actin dynamics via characterization of primitive gelsolin/cofilin proteins from Asgard archaea. Proc. Natl. Acad. Sci. USA 577, 202009167 (2020).

19. Leão, P. et al. Asgard archaea defense systems and their roles in the origin of eukaryotic immunity. Nat. Commun. 15, 6386 (2024).

20. Shomar, H. et al. Viperin immunity evolved across the tree of life through serial innovations on a conserved scaffold. *Nat*. Ecol. Evol. 8, 1667–1679 (2024).

21. Spang, A. et al. Proposal of the reverse flow model for the origin of the eukaryotic cell based on comparative analyses of Asgard archaeal metabolism. Nat. Microbiol. 4, 1138–1148 (2019).

22. Vosseberg, J. et al. Timing the origin of eukaryotic cellular complexity with ancient duplications. *Nat*. Ecol. Evol. 5, 92–100 (2020).

23. Stairs, C. W. & Ettema, T. J. G. The archaeal roots of the eukaryotic dynamic actin cytoskeleton. Curr. Biol. 30, R521–R526 (2020).

24. Sousa, F. L., Neukirchen, S., Allen, J. F., Lane, N. & Martin, W. F. Lokiarchaeon is hydrogen dependent. Nat. Microbiol. 1, 16034 (2016).

25. Seitz, K. W., Lazar, C. S., Hinrichs, K.-U., Teske, A. P. & Baker, B. J. Genomic reconstruction of a novel, deeply branched sediment archaeal phylum with pathways for acetogenesis and sulfur reduction. ISME J. 10, 1696–1705 (2016).

26. Bulzu, P.-A. et al. Casting light on Asgardarchaeota metabolism in a sunlit microoxic niche. Nat. Microbiol. 4, 1129–1137 (2019).

27. Appler, K. E. et al. Oxygen metabolism in descendants of the archaeal-eukaryotic ancestor. Preprint at https://www.biorxiv.org/content/10.1101/2024.07.04.601786v1 (2024).

28. Baughn, A. D. & Malamy, M. H. The strict anaerobe *Bacteroides fragilis* grows in and benefits from nanomolar concentrations of oxygen. Nature 427, 441–444 (2004).

29. Nobu, M. K. et al. Catabolism and interactions of uncultured organisms shaped by eco-thermodynamics in methanogenic bioprocesses. Microbiome 8, 111 (2020).

30. Lu, Z. & Imlay, J. A. When anaerobes encounter oxygen: mechanisms of oxygen toxicity, tolerance and defence. Nat. Rev. Microbiol. 19, 774–785 (2021).

31. Imachi, H. et al. Cultivation of previously uncultured microorganisms with a continuous-flow down-flow hanging sponge (DHS) bioreactor, using a syntrophic archaeon culture obtained from deep marine sediment as a case study. Nat. Protoc. 17, 2784–2814 (2022).

32. Zhu, Q.-Z., Wegener, G., Hinrichs, K.-U. & Elvert, M. Activity of ancillary heterotrophic community members in anaerobic methane-oxidizing cultures. Front. Microbiol. 13, 912299 (2022).

33. Urai, A. et al. Origin of deep methane associated with a unique community of microorganisms in an organic- and iodine-rich aquifer. ACS Earth Space Chem. 5, 1– 11 (2021).

34. Avcı, B. et al. Spatial separation of ribosomes and DNA in Asgard archaeal cells. ISME J. 16, 606–610 (2021).

35. Wolferen, M. van, Pulschen, A. A., Baum, B., Gribaldo, S. & Albers, S.-V. The cell biology of archaea. Nat. Microbiol. 7, 1744–1755 (2022).

36. Abramson, J. et al. Accurate structure prediction of biomolecular interactions with AlphaFold 3. Nature 630, 493–500 (2024).

37. Shimoyama, T., Kato, S., Ishii, S. & Watanabe, K. Flagellum mediates symbiosis. Science 323, 1574 (2009).

38. Lovley, D. R. Syntrophy goes electric: direct interspecies electron transfer. Annu. Rev. Microbiol. 71, 643–664 (2017).

39. Walker, D. J. F. et al. The archaellum of *Methanospirillum hungatei* is electrically conductive. mBio 10, e00579–19. (2019).

40. Eren, E., Watts, N. R., Montecinos, F. & Wingfield, P. T. Encapsulated ferritin-like proteins: a structural perspective. Biomolecules 14, 624 (2024).

41. Schink, B. & Stams, A. J. M. in The Prokaryotes: Prokaryotic Communities and Ecophysiology. (ed. Rosenberg, E. et al.) 471–493 (Springer, 2013).

42. Nakagawa, S. et al. Characterization of protein glycosylation in an Asgard archaeon. BBA Adv. 6, 100118 (2024).

43. Donachie, W. D., Begg, K. J. & Vicente, M. Cell length, cell growth and cell division. Nature 264, 328–333 (1976).

44. Sargent, M. G. Control of cell length in *Bacillus subtilis*. J. Bacteriol. 123, 7–19 (1975).

45. Preisig, O., Zufferey, R., Thöny-Meyer, L., Appleby, C. A. & Hennecke, H. A high-affinity *cbb*_3_-type cytochrome oxidase terminates the symbiosis-specific respiratory chain of *Bradyrhizobium japonicum*. J. Bacteriol. 178, 1532–1538 (1996).

46. Slobodkina, G. B., Lebedinsky, A. V., Chernyh, N. A., Bonch-Osmolovskaya, E. A. & Slobodkin, A. I. *Pyrobaculum ferrireducens* sp. nov., a hyperthermophilic Fe(III)-, selenate- and arsenate-reducing crenarchaeon isolated from a hot spring. Int. J. Syst. Evol. Microbiol. 65, 851–856 (2015).

47. Ames, B. N., Cathcart, R., Schwiers, E. & Hochstein, P. Uric acid provides an antioxidant defense in humans against oxidant- and radical-caused aging and cancer: a hypothesis. Proc. Natl. Acad. Sci. USA 78, 6858–6862 (1981).

48. Elek, A., Kuzman, M. & Vlahovicek, K. coRdon: Codon usage analysis and prediction of gene expressivity. R package version 1.24.0. https://www.bioconductor.org/packages/release/bioc/html/coRdon.html (2024).

49. Kaster, A.-K., Moll, J., Parey, K. & Thauer, R. K. Coupling of ferredoxin and heterodisulfide reduction via electron bifurcation in hydrogenotrophic methanogenic archaea. Proc. Natl. Acad. Sci. USA 108, 2981–2986 (2011).

50. Mei, R., Kaneko, M., Imachi, H. & Nobu, M. K. The origin and evolution of methanogenesis and *Archaea* are intertwined. PNAS Nexus pgad023 (2023).

## Methods references

51. Maekawa, T., Igari, S. & Kaneko, N. Chemical and isotopic compositions of brines from dissolved-in-water type natural gas fields in Chiba, Japan. Geochem. J. 40, 475 (2007).

52. Nakahara, N., et al. *Aggregatilinea lenta* gen. nov., sp. nov., a slow-growing, facultatively anaerobic bacterium isolated from subseafloor sediment, and proposal of the new order *Aggregatilineales* ord. nov. within the class *Anaerolineae* of the phylum *Chloroflexi*. Int. J. Syst. Evol. Microbiol. 69, 1185–1194 (2019).

53. Ludwig, W. et al. ARB: a software environment for sequence data. Nucleic Acids Res. 32, 1363–1371 (2004).

54. Quast, C. et al. The SILVA ribosomal RNA gene database project: improved data processing and web-based tools. Nucleic Acids Res. 41, D590–D596 (2013).

55. Nunoura, T. et al. Microbial diversity in deep-sea methane seep sediments presented by SSU rRNA gene tag sequencing. Microb. Environ. 27, 382–390 (2012).

56. Pfennig, N. & Wagener, S. An improved method of preparing wet mounts for photomicrographs of microorganisms. J. Microbiol. Methods 4, 303–306 (1986).

57. Cabral, B. & Leedom, L. C. Imaging vector fields using line integral convolution. Proc. 20th Annu. Conf. Comput. Graph. Interact. Tech. 263–270 (1993) doi:10.1145/166117.166151.

58. Paszke, A. et al. PyTorch: An imperative style, high-performance deep learning library. Proceedings of the 33rd International Conference on Neural Information Processing Systems Article No. 721, 8026–8037 (2019).

59. Fouard, C., Malandain, G., Prohaska, S. & Westerhoff, M. Blockwise processing applied to brain microvascular network study. IEEE Trans. Med. Imaging 25, 1319– 1328 (2006).

60. Tahara, Y. O. & Miyata, M. in Bacterial and Archaeal Motility: Visualization of peptidoglycan structures of Escherichia coli by quick-freeze deep-etch electron microscopy. (eds. Minamino, T., Miyata, M. & Namba, K.) Methods in Molecular Biology, vol. 2646, 299–307 (2023).

61. Kremer, J. R., Mastronarde, D. N. & McIntosh, J. R. Computer visualization of three-dimensional image data using IMOD. J. Struct. Biol. 116, 71–76 (1996).

62. Yamaguchi, T. et al. In situ DNA-hybridization chain reaction (HCR): a facilitated in situ HCR system for the detection of environmental microorganisms. Environ. Microbiol. 17, 2532–2541 (2015).

63. Toyooka, K. & Shinozaki-Narikawa, N. Efficient fluorescence recovery using antifade reagents in correlative light and electron microscopy. Microscopy 68, 417– 421 (2019).

64. Hirakata, Y. et al. Identification and cultivation of anaerobic bacterial scavengers of dead cells. ISME J. 17, 2279–2289 (2023).

65. Isaji, Y., Ogawa, N. O., Takano, Y. & Ohkouchi, N. Quantification and carbon and nitrogen isotopic measurements of heme B in environmental samples. Anal. Chem. 92, 11213–11222 (2020).

66. Maruo, M., Doi, T. & Obata, H. Onboard determination of submicromolar nitrate in seawater by anion-exchange chromatography with lithium chloride eluent. Anal. Sci. 22, 1175–1178 (2006).

67. Bolger, A. M., Lohse, M. & Usadel, B. Trimmomatic: a flexible trimmer for Illumina sequence data. Bioinformatics 30, 2114–2120 (2014).

68. Leggett, R. M., Clavijo, B. J., Clissold, L., Clark, M. D. & Caccamo, M. NextClip: an analysis and read preparation tool for Nextera Long Mate Pair libraries. Bioinformatics 30, 566–568 (2014).

69. Bankevich, A. et al. SPAdes: a new genome assembly algorithm and its applications to single-cell sequencing. J. Comput. Biol. 19, 455–477 (2012).

70. Sahlin, K., Chikhi, R. & Arvestad, L. Assembly scaffolding with PE-contaminated mate-pair libraries. Bioinformatics 32, 1925–1932 (2016).

71. Qin, M. et al. LRScaf: improving draft genomes using long noisy reads. BMC Genomics 20, 955 (2019).

72. Nadalin, F., Vezzi, F. & Policriti, A. GapFiller: a de novo assembly approach to fill the gap within paired reads. BMC Bioinformatics 13, S8 (2012).

73. Hyatt, D., LoCascio, P. F., Hauser, L. J. & Uberbacher, E. C. Gene and translation initiation site prediction in metagenomic sequences. Bioinformatics 28, 2223–2230 (2012).

74. Kalvari, I. et al. Rfam 13.0: shifting to a genome-centric resource for non-coding RNA families. Nucleic Acids Res. 46, D335–D342 (2017).

75. Lowe, T. M. & Chan, P. P. tRNAscan-SE On-line: integrating search and context for analysis of transfer RNA genes. Nucleic Acids Res. 44, W54–W57 (2016).

76. Langmead, B. & Salzberg, S. L. Fast gapped-read alignment with Bowtie 2. Nat. Methods 9, 357–359 (2012).

77. Liao, Y., Smyth, G. K. & Shi, W. featureCounts: an efficient general purpose program for assigning sequence reads to genomic features. Bioinformatics 30, 923–930 (2014).

78. Kawai, S., Shimamura, S., Shimane, Y. & Tsukatani, Y. Proteomic time-course analysis of the filamentous anoxygenic phototrophic bacterium, *Chloroflexus aurantiacus*, during the transition from respiration to phototrophy. Microorganisms 10, 1288 (2022).

79. Hashimoto, Y. et al. Physiological and comparative proteomic characterization of *Desulfolithobacter dissulfuricans* gen. nov., sp. nov., a novel mesophilic, sulfur-disproportionating chemolithoautotroph from a deep-sea hydrothermal vent. Front. Microbiol. 13, 1042116 (2022).

80. Nakagawa, S. et al. N-linked protein glycosylation in *Nanobdellati* (formerly DPANN) archaea and their hosts. J. Bacteriol. 206, e00205–24 (2024).

81. Parks, D. H. et al. GTDB: an ongoing census of bacterial and archaeal diversity through a phylogenetically consistent, rank normalized and complete genome-based taxonomy. Nucleic Acids Res. 50, gkab776 (2021).

82. Ondov, B. D. et al. Mash: fast genome and metagenome distance estimation using MinHash. Genome Biol. 17, 132 (2016).

83. Seemann, T. Prokka: rapid prokaryotic genome annotation. Bioinformatics 30, 2068– 2069 (2014).

84. Huerta-Cepas, J. et al. eggNOG 5.0: a hierarchical, functionally and phylogenetically annotated orthology resource based on 5090 organisms and 2502 viruses. Nucleic Acids Res. 47, D309–D314 (2019).

85. Minh, B. Q. et al. IQ-TREE 2: New models and efficient methods for phylogenetic inference in the genomic era. Mol. Biol. Evol. 37, 1530–1534 (2020).

86. Criscuolo, A. & Gribaldo, S. BMGE (Block Mapping and Gathering with Entropy): a new software for selection of phylogenetic informative regions from multiple sequence alignments. BMC Evol. Biol. 10, 210 (2010).

87. Nishihara, A., Tsukatani, Y., Azai, C. & Nobu, M. K. Illuminating the coevolution of photosynthesis and Bacteria. Proc. Natl. Acad. Sci. USA 121, e2322120121 (2024).

88. Katoh, K. & Standley, D. M. MAFFT Multiple sequence alignment software version 7: improvements in performance and usability. Mol. Biol. Evol. 30, 772–780 (2013).

89. Kalyaanamoorthy, S., Minh, B. Q., Wong, T. K. F., Haeseler, A. von & Jermiin, L. S. ModelFinder: fast model selection for accurate phylogenetic estimates. Nat. Methods 14, 587–589 (2017).

90. Lemoine, F. et al. Renewing Felsenstein’s phylogenetic bootstrap in the era of big data. Nature 556, 452–456 (2018).

91. Steinegger, M. & Söding, J. MMseqs2 enables sensitive protein sequence searching for the analysis of massive data sets. Nat. Biotechnol. 35, 1026–1028 (2017).

92. Price, M. N., Dehal, P. S. & Arkin, A. P. FastTree 2-approximately maximum-likelihood trees for large alignments. PLoS ONE 5, (2010).

93. Capella-Gutierrez, S., Silla-Martinez, J. M. & Gabaldon, T. trimAl: a tool for automated alignment trimming in large-scale phylogenetic analyses. Bioinformatics 25, 1972–1973 (2009).

94. Markin, A. et al. PARNAS: Objectively selecting the most representative taxa on a phylogeny. Syst. Biol. 72, 1052–1063 (2023).

95. Rozewicki, J., Li, S., Amada, K. M., Standley, D. M. & Katoh, K. MAFFT-DASH: integrated protein sequence and structural alignment. Nucleic Acids Res. 47, W5–W10 (2019).

96. Okuda, S. et al. jPOSTrepo: an international standard data repository for proteomes. Nucleic Acids Res. 45, D1107–D1111 (2017).

