## Extended Data Figures 1-10 for "Eukaryotes’ closest relatives are internally simple syntrophic archaea"

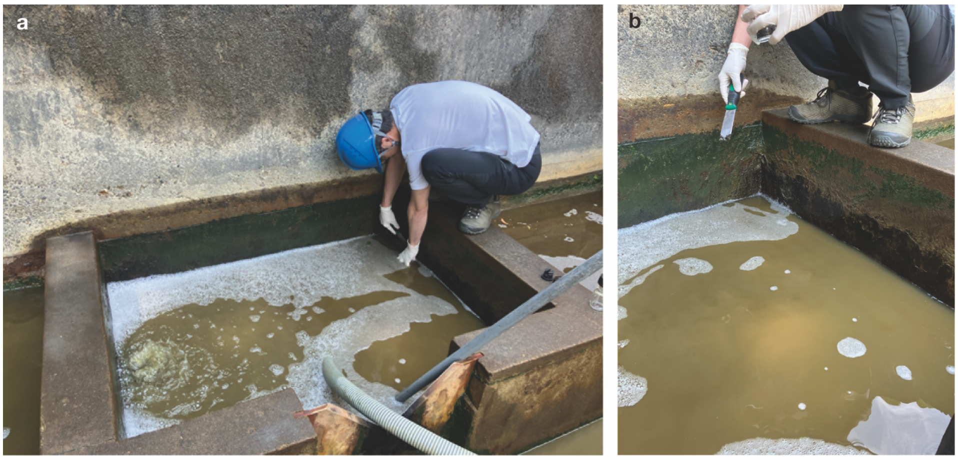

c

| Property | Unit | |
|---|---|---|
| $Na^+$ | mg/l | 7563 |
| $NH_4^+$ | mg/l | 167 |
| $K^+$ | mg/l | 223 |
| $Mg^{2+}$ | mg/l | 196 |
| $Ca^{2+}$ | mg/l | 104 |
| $Cl^-$ | mg/l | 12,669 |
| $Br^-$ | mg/l | 83.5 |
| $SO_4^{2-}$ | mg/l | Not Detected |
| $I^-$ | mg/l | 70.6 |
| Total Fe[a] | mg/l | 3.85 |
| Dissolved Fe[a] | mg/l | 0.23 |
| Total organic carbon (TOC) | mg/l | 69 |
| Oxygen Reduction Potential (ORP) | mV | -261 |
| Depth | m | 274–715 |
| Temperature | °C | 21.9 |
| Number of cells | cells/ml | $7.04\times10^7$ |

[a]These data were made public for the first time in this study.

This table is adapted from Supplementary Table S1 in Urai et al., 2021. ACS Earth and Space Chemistry, 5, 1–11., CC BY 4.0.

*c.f.*, Average Fe concentration in modern seawater is 0.03 μg/l (https://www.mbari.org/know-your-ocean/periodic-table-of-elements-in-the-ocean/).

**Extended Data Fig. 1 | Gas production well KTG3. a**, Biofilm sampling from the wall of the separator discharge outlet of the well. **b**, Biofilm on the wall. The water level in this well fluctuates every few hours due to the eruption of brine. The biofilm, which is normally in an anaerobic state, was sampled when the water level dropped after the eruption. **c**, Geochemical characteristics of brine in the well.

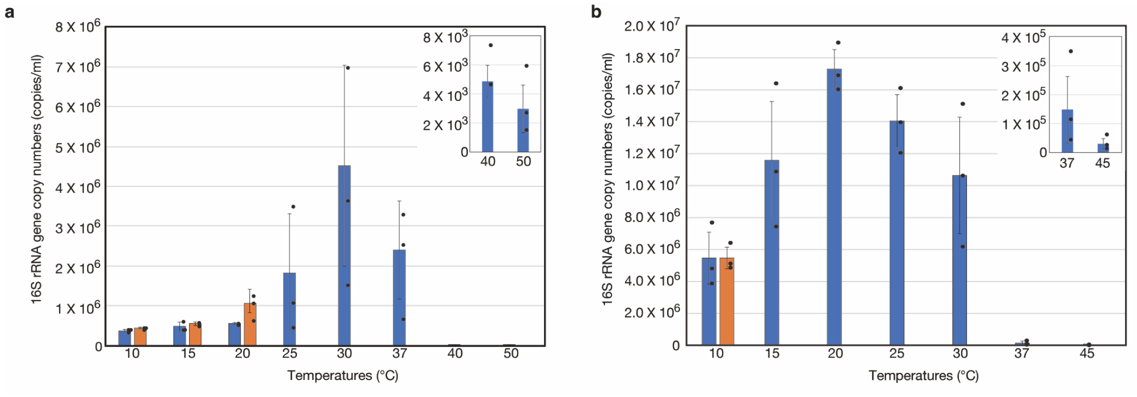


**Extended Data Fig. 2 | Effect of temperature on growth of cultured archaea. a**, HC1 cultures. Blue and orange bars show the 16S rRNA gene copy numbers after 32 and 62 days of incubation, respectively. Cultures incubated at 10°C, 15°C, and 20°C were monitored for a total of 62 days and then analyzed by qPCR. **b**, SC1 cultures. Blue and orange bars show the 16S rRNA gene copy numbers after 57 and 113 days of incubation, respectively. Cultures incubated at 10°C were monitored for a total of 113 days and then analyzed by qPCR. Data are mean ± s.d. of triplicate determinations. Each data point is shown as a dot.

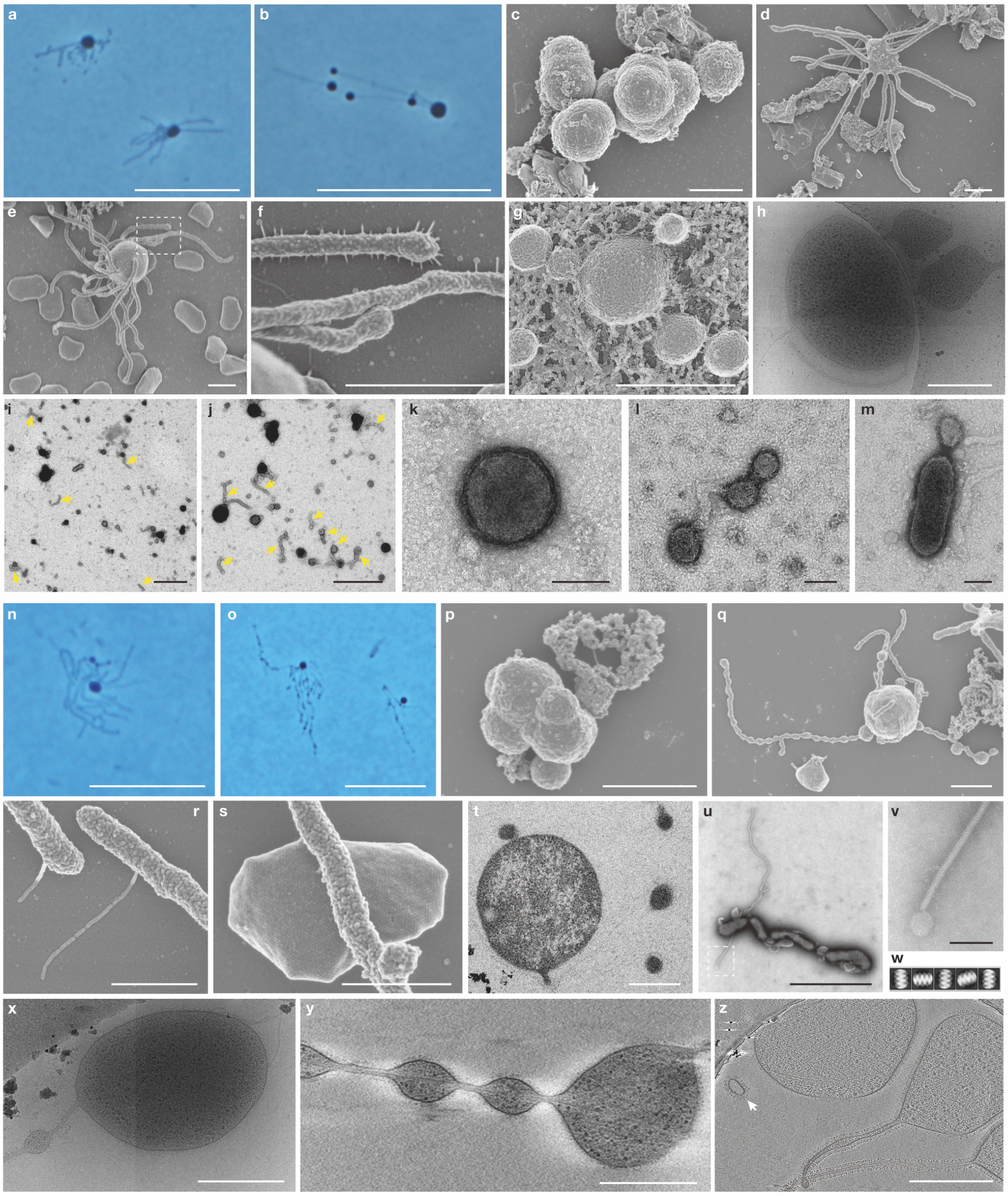


**Extended Data Fig. 3 | Other representative photomicrographs of HC1 and SC1. a–m**, HC1 cultures. **a**, **b**, Phase-contrast micrographs of unfixed cells. **c–f**, SEM images showing aggregated (**c**), protrusion-producing (**d**), and methanogen-attached (**e**) cells and short filaments on the surface of protrusions (**f**). The image **f** is a magnified view of the dashed box in (**e**). **g**, QFDE image of HC1. **h,** Slice image from a cryo-tomogram of an HC1 cell cultured with 0.05% yeast extract. **i–m**, TEM images of negatively stained membrane vesicles obtained from HC1 cultures. The yellow arrows in images **i** and **j**

indicate filamentous structures that appear to have detached from the protrusions. **n**–**z**, SC1 cultures. **n**, **o**, Phase-contrast micrographs of unfixed SC1 cells. **p**–**s**, SEM images of SC1 cultures, showing aggregated cells (**p**), a cell producing protrusions with bulging sections (**q**), short filaments on the surface of protrusions (**r**), and a protrusion adhering to a partner methanogen cell vis a short filament (**s**). The image **r** is a magnified view corresponding to Fig. 2q. **t**, Ultrathin section of an SC1 cell containing a vacuole or an internal vesicle. **u, v**, TEM images of negatively stained protrusion and thin filaments from unfixed sample. The image **v** shows a magnified view of the dashed box in **u**. **w**, Two-dimensional averaged images of the thin filament. **x**–**z**, Cryo-electron tomography images of SC1. The white arrow in image **z** indicates a membrane vesicle. The images **a**, **b**, **d**, **g**–**q**, **t**, **x**–**z** were obtained from cultures in the late-exponential phase, and the others were obtained from cultures in the mid-exponential phase. The phase-contrast microscopic images of HC1 and SC1 are representatives of $n$ = 126 and 65 recorded images, respectively. The TEM images of negatively stained HC1 membrane vesicles are representative of $n$ = 29 recorded images. The TEM image of negatively stained SC1 protrusion and thin filaments represent $n$ = 2 recorded images. Scale bars, 10 µm (**a**, **b**, **n**, **o**), 1 µm (**c**–**g**, **i**, **j**, **p**, **q**, b), 500 nm (**h**, **r**–**t**, **x**–**z**), and 100 nm (**k**–**m**, **v**).

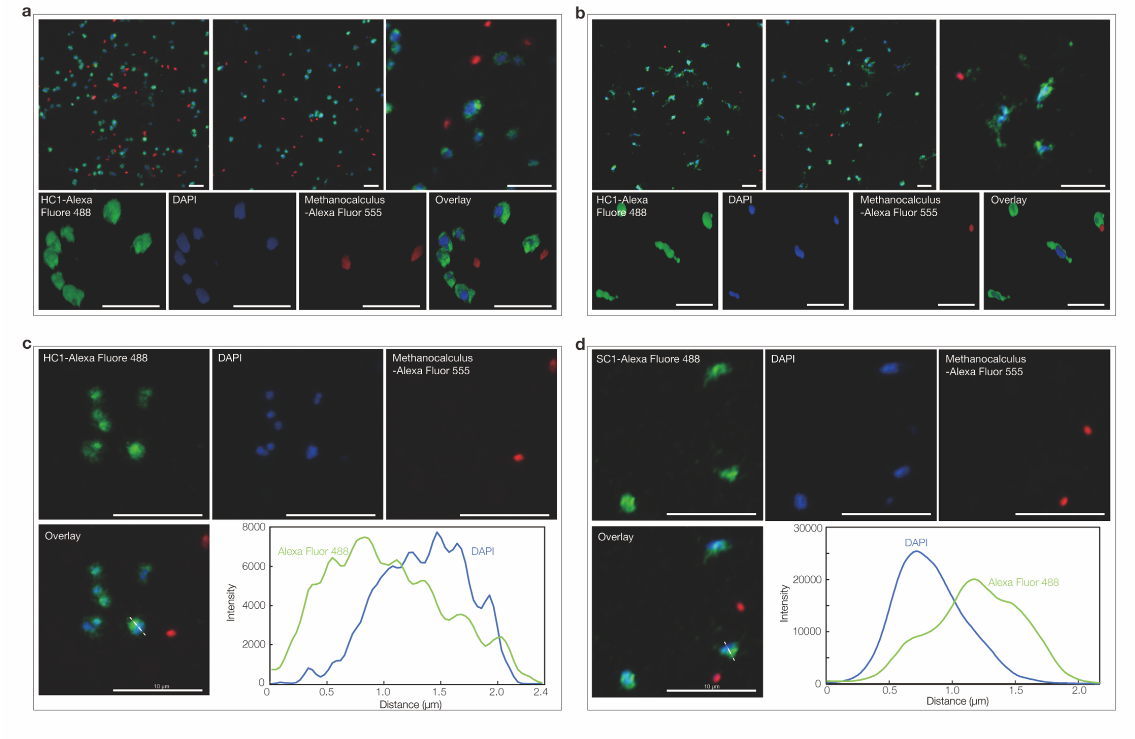


**Extended Data Fig. 4 | Other representative HCR-FISH images of cultured archaea.** **a**, Fluorescence images of cells from a pure co-culture of HC1 and *Methanocalculus* MC2 stained with DAPI (blue) and hybridized with nucleotide probes that target HC1 cells (green) and *Methanocalculus* (red). Four small panels at the bottom show a single three-dimensional (3D) image of each fluorescence and their merged images. A 360° rotation of the 3D image is also provided in Supplementary Video 2. **b**, Fluorescence images of cells from a tri-culture of SC1, *Methanocalculus* MC3 and *Methanolobus* MBL stained with DAPI (blue) and hybridized with nucleotide probes that target SC1 cells (green) and *Methanocalculus* (red). Four small panels at the bottom show a single 3D image of each fluorescence and their merged images. A 360° rotation of the 3D image is also provided in Supplementary Video 2. **c**, **d**, fluorescence intensity line profile of FISH probe and DAPI signals in HCR-FISH images of HC1 (**c**) and SC1 (**d**) cultures. Dashed lines indicate the position at which the fluorescence intensity line profiles were recorded. The red signals are not shown in the profiles because no red signals were derived from *Methanocalculus* MC2 cells at the locations where the intensity profiles were recorded. All the FISH images were acquired by the confocal laser scanning microscopy with a high-resolution mode (*i.e.*, Lightning technology of Leica). Scale bars, 5 µm (**a**, **b**) and 10 µm (**c**, **d**).

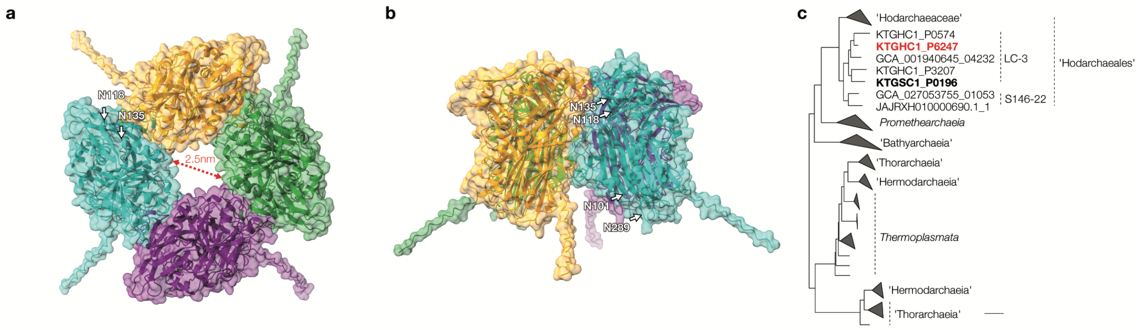


**Extended Data Fig. 5 | Predicted structure and phylogeny of the HC1 S-layer protein. a–b**, Side-view (**a**) and top-view (**b**) of the structure of the putative HC1 S-layer protein predicted using AlphaFold3 (KTGHC1_P6247). Asparagine residues detected to be N-glycosylated are indicated with arrows. The diameter of the central vertical pore is shown. **c**, Maximum likelihood estimation of the phylogeny of the putative S-layer protein and homologs detected in other archaea. All branches with ultrafast bootstrap support lower than 0.95 are collapsed.

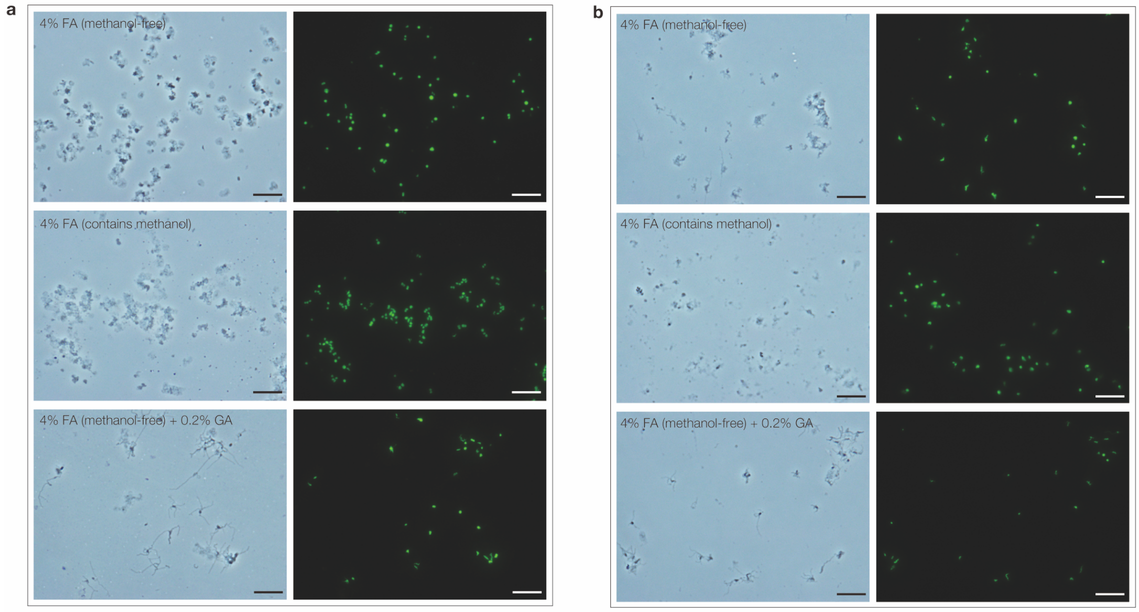


**Extended Data Fig. 6 | Microscopy images of chemically fixed cells of HC1 and SC1.** **a**, Fixed cells of HC1 culture. **b**, Fixed cells of SC1 culture. Fixing solution names (formaldehyde [FA], and glutaraldehyde [GA]) are indicated for each panel. The fluorescence images on the right side show cells stained with SYBR Green I. Each fluorescence image is in the same field as the image on the left. Prior to the fixation, we confirmed that HC1 and SC1 cells produced many protrusions in these cultures using phase contrast microscopy. Scale bars, 10 μm.

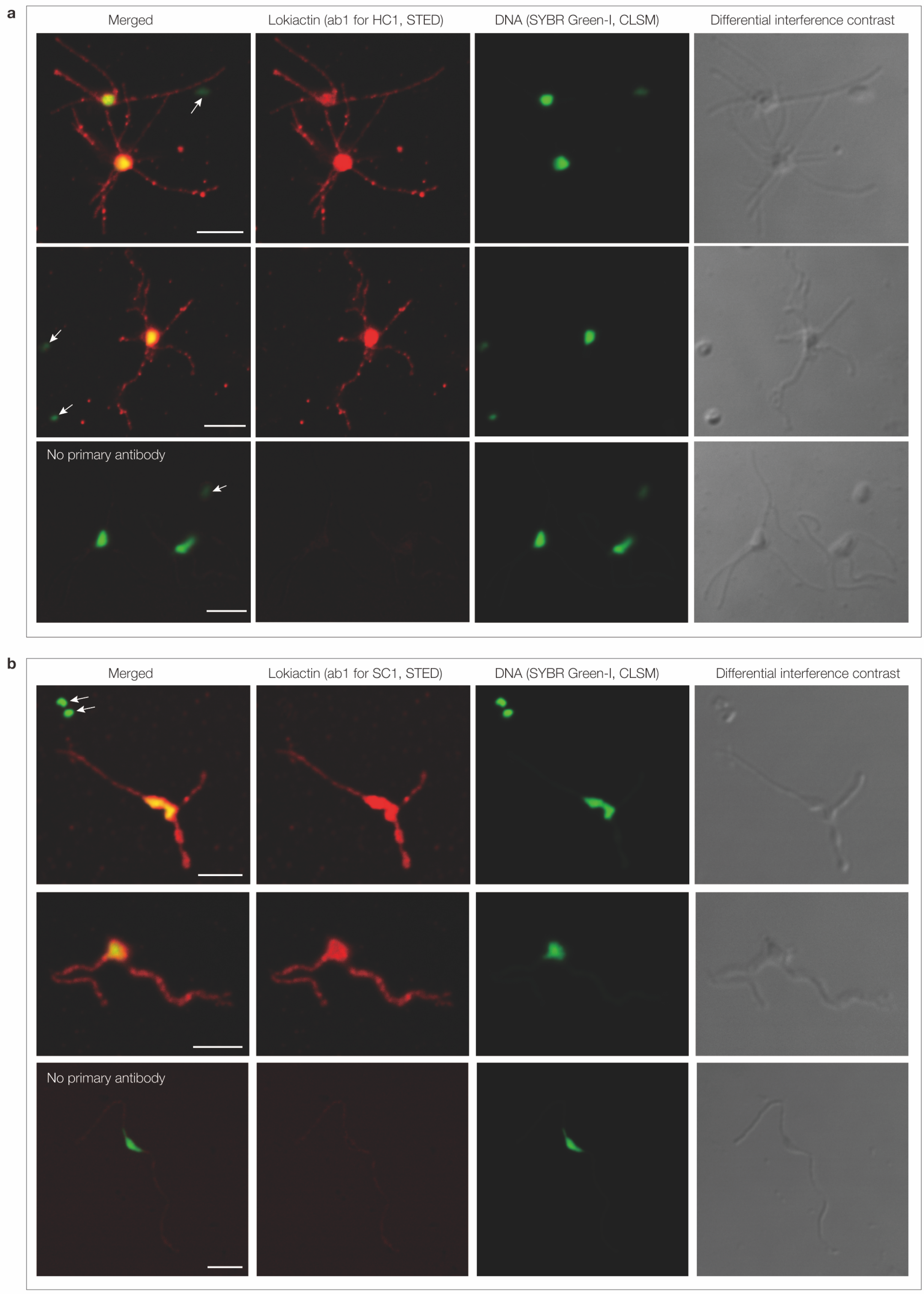


**Extended Data Fig. 7 | Immunofluorescence staining of HC1 and SC1 cells with lokiactin-specific antibodies imaged using a combination of confocal laser scanning microscopy (CLSM) and stimulated emission depletion (STED). a**, HC1 culture. **b**, SC1 culture. The detection of the lokiactin and DNA are shown in red (abberior

STAR580-labeled secondary antibody) and yellow/green (SYBR Green-I), respectively. White arrows indicate *Methanocalculus* cells. Scale bars, 3 µm. Note that some merged images are the same as those shown in Fig. 2j and x.

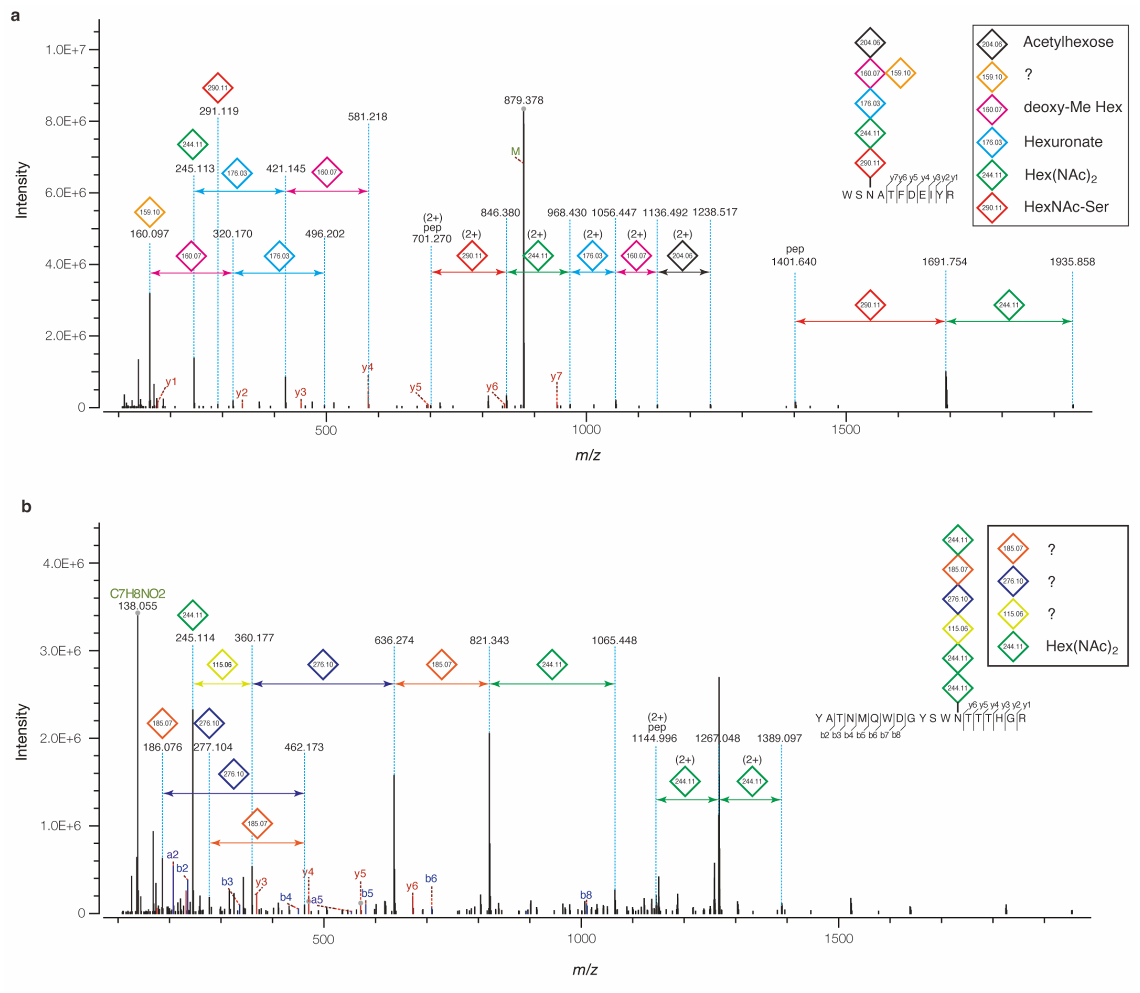


**Extended Data Fig. 8 | MS/MS spectra showing glycan assignments on proteins, including probable monosaccharides symbolized with their molecular weights. a,** Higher-energy collision dissociation (HCD) spectrum showing the *N*-glycan structure on peptide/nickel transport system substrate-binding protein (KTGHC1_P1252) of strain HC1 grown on casamino acids, peptone and yeast extract (0.005%, 0.005%, 0.05%, respectively). **b**, HCD spectrum showing the *N*-glycan on peptide/nickel transport system substrate-binding protein (KTGSC1_P0547) of strain SC1.

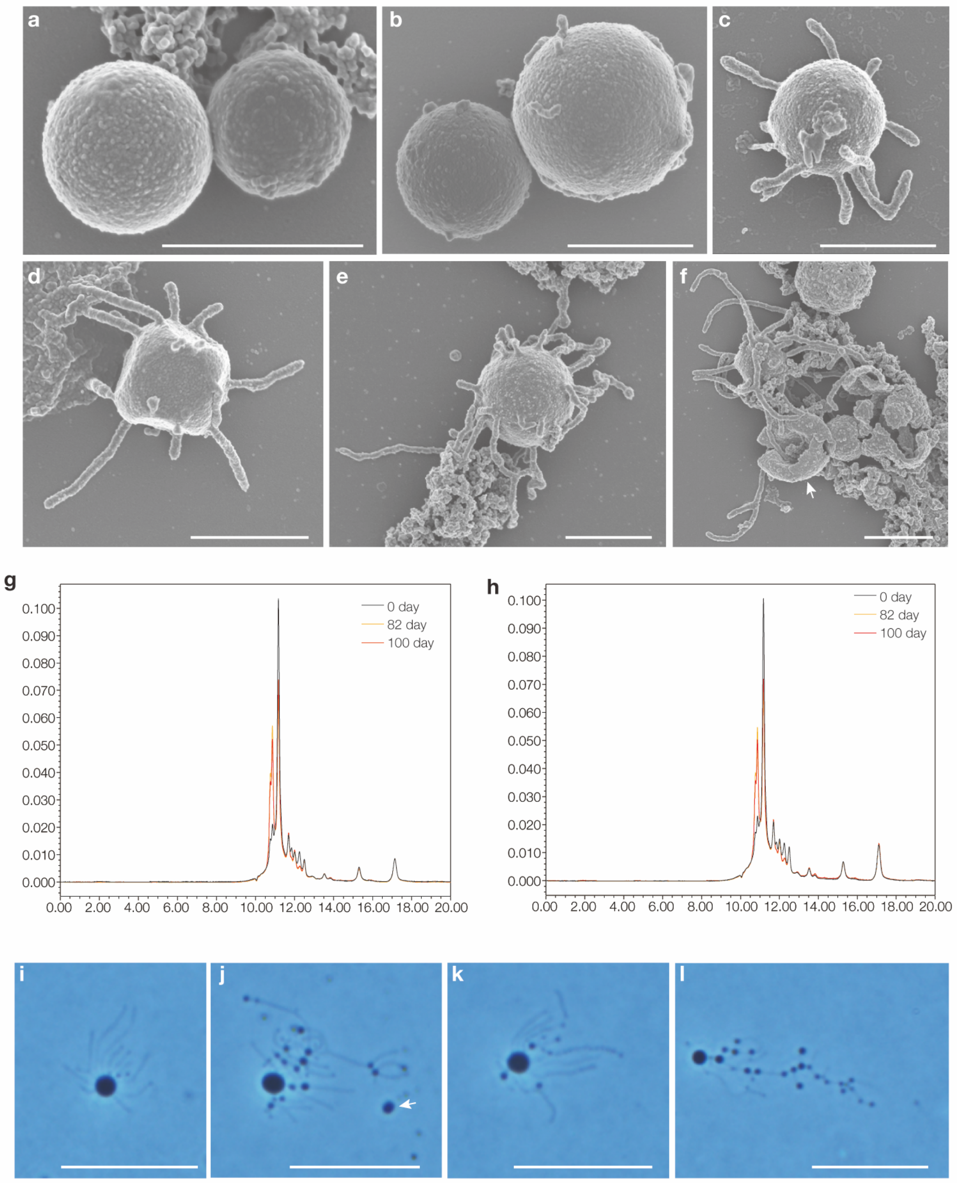


**Extended Data Fig. 9 | *P. syntrophicum* strain MK-D1 and strain SC1 grown in medium containing high concentration of yeast extract. a-f**, SEM images of *P. syntrophicum* MK-D1 cultured with yeast extract (0.05%, w/v). **g**, **h**, Changes in high molecular weight compounds in the culture medium under the same conditions. **i-l**, Phase contrast micrographs of SC1 cultured with 0.5% yeast extract, sampled 43 days after the start of incubation. In addition to yeast extract, the medium also contained casamino acids, powdered milk, and polypeptone peptone. White arrows indicate their partner methanogens (*i.e.*, *Methanogenium* sp. MK-MG and *Methanocalculus* sp. MC3). Scale bars, 1 µm (**a-f**) and 10 µm (**i-l**).

**a**

| Culture ID | Amount of nitrate reduced (μM) | Amount of nitrite produced (μM) | Methane concentration (μM) | Percentage of electrons used for nitrate-reduction (%)[a] | 16S rRNA gene copy numbers (copies/ml) | DNA concentration (ng/μl)[b] |
|---|---|---|---|---|---|---|
| ***1st time cultivation experiment with approximately 100 μM nitrate (incubation for 26 days)*** | | | | | | |
| No.1 Casamino acids + Peptone + Yeast extract w/o nitrate (control) | - | - | Not analyzed | - | 3.32E+06 | 1.44 |
| No.2 Casamino acids + Peptone + Yeast extract + nitrate | 10.1 | 11.2 | 19.2 | 11.6 | 2.29E+06 | 1.73 |
| No.3 Casamino acids + Peptone + Yeast extract + nitrate | 12.3 | 11.3 | 19.2 | 13.8 | 2.93E+06 | 1.86 |
| ***2nd time cultivation experiment with approximately 100 μM nitrate (incubation for 27 days)*** | | | | | | |
| No.1 Casamino acids + Peptone + Yeast extract w/o nitrate (control) | - | - | 30.9 | - | 5.33E+06 | 1.88 |
| No.2 Casamino acids + Peptone + Yeast extract w/o nitrate (control) | - | - | 32.7 | - | 5.15E+06 | 1.95 |
| No.3 Casamino acids + Peptone + Yeast extract + nitrate | 19.2 | 19.9 | 26.8 | 15.2 | 3.80E+06 | 2.1 |
| No.4 Casamino acids + Peptone + Yeast extract + nitrate | 16.8 | 16.1 | 26.9 | 13.5 | 3.88E+06 | 3.08 |
| No.5 Casamino acids + Peptone + Yeast extract + nitrate + 2-BES[c] | 0 | ND | ND | - | 1.94E+05 | 0.124 |
| No.6 Casamino acids + Peptone + Yeast extract + nitrate + 2-BES | 0 | ND | ND | - | 1.32E+03 | 0.16 |
| No.7 Casamino acids + Peptone + Yeast extract + nitrate w/o reducing agents | 0 | ND | ND | - | 6.24E+04 | ND |
| No.8 Casamino acids + Peptone + Yeast extract + nitrate w/o reducing agents | 0 | ND | ND | - | 3.25E+04 | ND |

[a]Calculated based on the amount od electrons used for nitrate-reduction and the amount of electron used for methane production.

[b]DNAs were extracted from 1 ml of culture liquid and dissolved in 21 μL TE buffer. The DNA concentrations were quantified by Quant-iT dsDNA High-Sensitivity Assay Kit (Qiagen).

[c]2-bromoethanesulfonate (2-BES), a methanogenesis inhibitor, was added at a final concentration of 10 mM.

**b**

| Culture ID[a] | HC1 16S rRNA gene copies per ml of culture after 26 days of incubation | Final DNA concentration (ng/μl)[b] |
|---|---|---|
| No.1 Casamino acids + Peptone + Yest extract | 6.79E+06 | 2.2 |
| No.2 Casamino acids + Peptone + Yest extract | 5.06E+06 | 1.73 |
| No.3 Casamino acids + Peptone + Yest extract + Nitrate (1 mM) | 5.92E+04 | ND |
| No.4 Casamino acids + Peptone + Yest extract + Nitrate (1 mM) | 5.16E+04 | ND |

[a]The same parent culture was used for all the cultures as the inoculum.
[b]DNAs were extracted from 1 ml of culture liquid and dissolved in 21 μL TE buffer. The DNA concentrations were quantified by Quant-iT dsDNA High-Sensitivity Assay Kit (Qiagen).

**c**

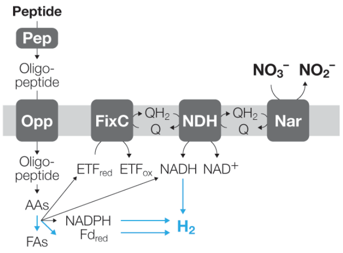


**Extended Data Fig. 10 | Nitrate utilization by HC1. a**, Cultivation tests of HC1 with addition of 100 μM sodium nitrate. **b**, Cultivation tests of HC1 with addition of 1 mM sodium nitrate. **c**, Schematic diagram of HC1 nitrate metabolism via nitrate reductase (Nar). See Figure 4 for explanation of abbreviations.
