## Supplemental Data 1 for "Eukaryotes’ closest relatives are internally simple syntrophic archaea"

**Supplementary Information**

This file contains Supplementary Notes 1–8, Supplementary Methods, Supplementary Figures 1–11, and Supplementary References.

**Table of Contents**

Supplementary Notes………02
Supplementary Methods……09
Supplementary Figures……..12
Supplementary References…24

### Supplementary Note 1

### The cultivation conditions we devised for culturing new members of *Promethearchaeota*.

As mentioned in the main text, we hypothesized that the sampled biofilm would serve as an ideal source for cultivating 'Heimdallarchaeia' and elucidating their relationship with oxygen. To cultivate them, we used our hypotheses and the following information to guide the establishment of specific cultivation conditions.

1. Indirect provision of energy substrates in primary enrichment cultures

   Previous studies have shown that *Promethearchaeota* could be co-enriched with microbial communities performing anaerobic oxidation of methane (AOM) and propionate (AOP)[1–3]. We hypothesized that members of *Promethearchaeota* can utilize organic matter (probably mainly amino acids/peptides) secreted by microorganisms that consume methane or propionate[2,4].

2. Use proteinaceous substrates to further enrich *Promethearchaeota*

   Comparative genomics showed that most members of *Promethearchaeota* are capable of anaerobic amino acids/peptides degradation[5]. Therefore, once we confirmed their co-enrichment in cultures amended with methane or propionates, we shifted to proteinaceous substrates (*e.g.*, casamino acids, powdered milk, peptone, and yeast extract) to promote further enrichment.

3. Cultivation at low concentrations of proteinaceous substrates

   *P. syntrophicum* strain MK-D1 is a slow-growing microorganism with extremely low cell yield[5,6]. Because many slow-growing microorganisms are known to exhibit high affinities for their energy substrates[7,8], we hypothesized members of *Promethearchaeota* would similarly possess a high substrate affinity for amino acids and peptides. Therefore, during the initial enrichment phase, we prepared a medium supplemented with low concentrations of amino acids/peptide compounds (0.005%, w/v). Note that *P. syntrophicum* strain MK-D1 could grow at this concentration of casamino acids and 20 amino acids, but scarcely at 0.0005% (w/v)[5].

4. Addition of antibiotics to enrich archaeal species

   We previously employed four antibiotics (i.e., ampicillin, vancomycin, kanamycin, and streptomycin) to enrich *P. syntrophicum* strain MK-D1[5]. In this study, we applied these and other antibiotics to enrich archaeal species more effectively. For example, erythromycin may serve as a promising candidate because a comparative genomic analysis targeting transporters[9] suggests that *Promethearchaeota* may tolerate it. Ciprofloxacin, azithromycin, cephalothin, and ofloxacin could also act as selective agents for enriching diverse archaeal species, as these antibiotics have been successfully used to enrich ammonium-oxidizing archaea, haloarchaea and methanogens[10–15].

5. Using natural brine as the basal medium for initial enrichment cultures
   Establishing cultivation conditions that closely resemble the natural environment is a fundamental strategy for culturing indigenous microorganisms. To maintain conditions as close to *in situ* as possible, we used brine (*i.e.*, gas-associated formation water) obtained anoxically from the gas-production well as the basal medium for the initial enrichment stage. After confirming stable growth of the target archaeon, we shifted to an artificial medium for further enrichment.
6. Adjusting reducing agent concentrations
   Some 'Heimdallarchaeia' possess terminal oxidases[16–20]. Though whether the enzymes support aerotolerance or aerobic respiration remains unclear, 'Heimdallarchaeia' likely have the capacity to help them survive and possibly grow in environments exposed to oxygen. To selectively cultivate these archaea, we either decreased or omitted reducing agents in the culture medium.

### Supplementary Note 2

**Cultivation of novel 'Hodarchaeales' strain SC1**. Since HC1 was confirmed to grow on artificial medium, we expected that other members of *Promethearchaeota* could also be enriched without relying on brine-based medium during the initial enrichment stage. We therefore directly inoculated fresh biofilm sample into artificial media on-site and incubated the culture vials at 20°C, a temperature near that of the sampling site brine (Extended Data Table 1).

After four months of incubation, 16S rRNA gene sequences affiliated with *Promethearchaeales* were detected in an anaerobic medium containing propionate and four antibiotics (*i.e.*, ampicillin, vancomycin, kanamycin, and streptomycin), although their relative abundance was less than 0.04% (Supplementary Table 1). We subsequently repeated subcultures in using the same medium composition twice and additionally detected a small number of sequences belonging to 'Hodarchaeales' (tentatively named HC2). To further enrich these archaea, we transferred them to a medium supplemented with casamino acids (0.005%, w/v), powdered milk (0.005%, w/v), and five antibiotics. After four months of incubation, a new 'Hodarchaeales' population (later strain SC1) appeared, with a relative abundance of 0.55%. This sequence had not been detected in the mother cultures, suggesting that the archaeon was initially a very minor population (Supplementary Table 1).

To further enrich SC1, we continued subcultures in the casamino acids-powdered milk-medium supplemented with the five antibiotics and monitored its growth by qPCR using SC1-specific 16S rRNA gene primers, as well as by iTag analysis of the community composition. During the enrichment process, we also tested three additional antibiotics (i.e., ciprofloxacin, cephalothin, and ofloxacin), to which HC1 was found to be resistant. After 182 days of incubation, SC1 reached a relative abundance of 23% with $10^5$ 16S rRNA gen copies $ml^{-1}$ in an anaerobic medium amended with casamino acids, powdered milk, the five antibiotics, and cephalothin (Supplementary Table 1).

To further increase the abundance, growth rate, and cell density of SC1, we added peptone and yeast extract to the medium as additional energy sources and tested the combination of the energy substrates/antibiotics, their concentrations, and cultivation temperatures. After three successive transfers with modified cultivation conditions, we obtained a tri-culture constituting of the target archaeon SC1, the hydrogenotrophic methanogen *Methanocalculus* sp. strain MC3, and the methylotrophic methanogen *Methanolobus* sp. strain MLB. This tri-culture has stably been maintained in MK medium amended with casamino acids (0.005%, w/v), powdered milk (0.005%), polypeptone peptone (0.01%, w/v), and yeast extract (0.05%, w/v) at 20°C (Supplementary Table 1). Under these cultivation conditions, SC1 required approximately 60 days to reach maximum growth with a cell density of $10^7$ 16S rRNA gene copies $ml^{-1}$ (Fig. 1e). Overall, the time from sampling to the establishment of the tri-culture was approximately three years.

**Supplementary Note 3**

**Taxonomic description for archaeal strains cultured in this study.**

**Etymology.** HC1: *Margulisarchaeum*, *Margulis* (Latin): named after Lynn Margulis, an evolutionary biologist; *archaeum* from *archaea* (Greek): an ancient life. The genus name is after Lynn Margulis, who proposed the endosymbiotic theory, a revolutionary idea in the origin of eukaryotes. The species name, *peptidophila*, *peptidum* (Latin): peptide; *phila* (Latin) loving. The species name refers to the peptide utilization property of HC1.

SC1: *Flexarchaeum*, *flexus* (Latin): flexible; *archaeum* from *archaea* (Greek): an ancient life. The genus name is derived from its flexible cells. The species name, *multiprotrusionis*, *multus* (Latin): many, multi; *protrusio* (Greek) protrusion. The species name is derived from the property of SC1 cells that produce many protrusions.

**Locality.** Both strains were obtained from biofilms formed on the wall of a gas production well, Chiba Prefecture, Japan.

**Diagnosis.** Anaerobic, peptides-degrading archaea. Coccoid-shaped cells. Syntrophically grows with hydrogen- and formate-utilizing microorganisms. HC1 reduces nitrate to nitrite. They produce membrane vesicles and membrane-based protrusions.

### Supplementary Note 4

### Peptide quantification

We used an UPLC-based analysis to quantify the consumption of macromolecules in the culture media and, as another approach, we tested the Pierce Quantitative Fluorometric Peptide Assay kit (Thermo Fisher Scientific). However, due to inconsistent measurement reproducibility, we discontinued its use. According to the manufacturer's instructions, amino acid-containing buffers are incompatible with this assay, suggesting that the cysteine employed as a reducing agent in the culture media may have interfered with the measurements.

### Supplementary Note 5

### Glycoproteome analysis of strains HC1 and SC1

*Protein diversity*

Our glycoproteome analysis using *N*-glycosylation open search identified 786 and 926 different proteins from strain HC1 grown on 0.005% and 0.05% yeast extract, respectively, representing approximately 12.0% and 14.1% of its total coding sequence (CDS). For strain SC1, 1,436 different proteins were identified, constituting about 26.2% of its total CDS. These proportions, similar to another member, *P. syntrophicum* MK-D1, where 19.1% of CDS was detected[21], are significantly lower compared to the approximately 50% previously reported for *Thermoproteota* archaea in our earlier analysis[22]. The lower proportion of detected proteins in *Promethearchaeota* archaea may be attributed to their larger genome sizes and higher gene numbers, which may result from increased gene duplications and reduced gene losses, unlike other archaeal lineages[17].

Throughout the two cultivation conditions, high normalized spectral abundance factors (NSAF) values were observed in HC1 for hypothetical proteins (KTGHC1_P0390 and KTGHC1_P6317), a trehalose/maltose transport system substrate-binding protein (KTGHC1_P2505), peptide/nickel transport system substrate-binding proteins (KTGHC1_P4283 and KTGHC1_P5849), and DNA-binding protein (KTGHC1_P0859). Similarly, in SC1, peptide/nickel transport system proteins (KTGSC1_P0547 and KTGSC1_P0136) exhibited high NSAF values, along with other hypothetical proteins. Similar substrate-binding proteins (SBPs) were abundantly detected in *P. syntrophicum*

MK-D1[21], suggesting the common significance of these SBPs in the nutritional strategies of *Promethearchaeota* archaea.

*Protein N-glycosylation*

The structure of archaeal *N*-glycans, including the composition and sequence of monosaccharides, provides important insights into the phylogenetic relationships among archaeal species. Protein *N*-glycosylation of HC1 and SC1 was examined as previously described[21,22]. Strain HC1 revealed a single predominant modification value of 1233.48 Da in both cultivation conditions. In contrast, modifications at 1308.55 Da were abundant in strain SC1, suggesting a different glycan structure. Many glycoproteins, including substrate-binding proteins and hypothetical proteins, were identified (Supplementary Table 7), all undergoing *N*-glycosylation at asparagine residues within the conserved N-X-S/T sequon. This glycosylation pattern, where X represents any amino acid except proline, is common across archaea, bacteria, and eukaryotes. Similar to previously reported archaeal S-layer proteins, S-layer proteins of HC1 (KTGHC1_P6247) and SC1 (KTGSC1_P0196) were *N*-glycosylated, primarily in the vicinity of the N-terminus (Supplementary Table 7 and Extended Data Fig. 8).

The MS/MS spectra of the glycopeptides from HC1 consistently identified unusual potential oxonium ions of sugars at 160.10, 245.11, and 291.12 Da. The MS/MS analysis revealed a probably linear glycan consisting of 290.11-244.11-176.03-160.07-159.10-204.06 (Extended Data Fig. 8a). Archaeal protein *N*-glycosylation is known for its diverse array of linking sugars[23]. An unusual linking sugar with an m/z of 290.11 Da, potentially HexNAc modified with serine (theoretical mass, 290.1114 Da), was previously reported as the linking sugar in the hyperthermophilic archaeon *Pyrodictium abyssi*, although the possibility of it being a different atypical sugar cannot be excluded[21]. Additionally, 244.11 Da mass may represent $Hex(NAc)_2$, as reported in the linking sugar of *Pyrobaculum calidifontis*[24]. The identification of two HexNAc variants at the reducing end suggests that the N-glycan of HC1 possesses a chitobiose core, similar to those found in *Thermoproteota* archaea and eukaryotes[23]. The masses of 176.03 Da and 160.07 Da are potentially uronic acid and deoxy-methyl hexose, respectively. Uronic acids and methylated sugars were previously reported in *N*-glycans of various *Methanobacteriota* archaea[23]. The 204.06 Da mass may correspond to acetyl-hexose. No candidate structure was identified for the mass of 159.10 Da, suggesting it may be an unknown component. In summary, the glycan structure of HC1 exhibits reducing ends characteristic of *Thermoproteota* and eukaryote glycans, middle parts of *Methanobacteriota* glycans, and unique non-reducing ends with unknown sugar components.

Similarly, the MS/MS spectra of the glycopeptides from SC1 consistently identified unusual potential oxonium ions of sugars at 186.08, 245.11, and 277.10 Da. The MS/MS analysis revealed a potentially linear glycan consisting of 244.11-244.11-115.06-276.10-185.07-244.22 (Extended Data Fig. 8b). As the glycan detected in HC1, the 244.11 Da mass may represent $Hex(NAc)_2$. No candidate structure was identified for the mass of 115.06, 185.07, and 276.10 Da, suggesting they may be unknown components. The glycan structure of strain SC1 features reducing ends characteristic of *Thermoproteota* and eukaryote glycans, with middle parts and non-reducing ends that include unknown sugar components. Previous glycoproteomic analysis identified a glycan structure in *P. syntrophicum* that lacks a chitobiose core[21].

In summary, glycan structure similarities were observed between HC1 and SC1, and the reducing end contained a structure shared with eukaryotes and *Thermoproteota* (a group of archaea most closely related to the *Promethearchaeota*). Although a previous study has reported that *P. syntrophicum* MK-D1 possesses glycan structures that are significantly different from those of eukaryotes[21], the current study suggests that *P. syntrophicum* MK-D1 may be an exceptional case and glycan structural similarities have been confirmed between eukaryotes and 'Hodarchaeales'.

**Supplementary Note 6**

*P. syntrophicum* MK-D1 and SC1 demonstrated behavior similar to HC1 when cultured at higher peptide concentrations. For MK-D1, when the substrate 0.05% casamino acids was replaced with 0.05% yeast extract, the doubling time shortened only slightly (from 14 days to 11 days)[6]. Meanwhile, the cell size nearly doubled (from 0.3–0.75 μm to 0.7–1.5 μm) and the number of protrusions per cell increased substantially (from a maximum of 8 to 14) (Extended Data Fig. 9). As for SC1, when the yeast extract concentration was raised to 0.5%, its cell size increased by approximately 1.5 times while the cell yield remained unchanged (Extended Data Fig. 9 and Supplementary Table 10).

**Supplementary Note 7**

Enzymes related to anaerobic peptidotrophy (extracellular peptidases, glutamate dehydrogenase, glycine dehydrogenase, 2-oxoacid:ferredoxin oxidoreductases, and hydrogenases) had codon utilization similar to ribosomal proteins, putting them in the top 25% among HC1's proteins (Fig. 3d; Supplementary Table 5). Supporting the validity of the approach, transcriptomics and proteomics both confirm that all the above genes/proteins were highly expressed by HC1 (Supplementary Table 5). Meanwhile, genes encoding cytochrome *c* oxidase, cytochrome *bd* oxidase-like protein, complex III cytochrome *b* (without cytochrome *c*/*f* or electron carrier copper proteins), type II NADH:quinone oxidoreductase, and nitrate reductase were all predicted to have low expressivity. Transcriptomics and proteomics detected low expression of these genes/proteins in anaerobic culture conditions despite the absence of $O_2$ and nitrate. This is in line with our inference that the genes likely support anaerobic peptidotrophy with aerotolerance.

**Supplementary Note 8**

The slow growth rate conserved among all cultured *Promethearchaeota* members could have theoretically played a critical role in the rise of eukaryotes. More specifically, we suspect the property may have contributed to closing several major gaps between FECA/archaea and eukaryotes: (i) absence/presence of a mitochondrion (endosymbiont), (ii) membrane lipid composition (ether vs ester lipids[25,26]), (iii) cell size, and (iv) genome composition. Firstly, assuming FECA somehow endogenized the protomitochondrion, for FECA to retain the endosymbiont, it must have had a growth rate slower than the protomitochondrion. In addition, the protomitochondrion very likely possessed the ability to respire oxygen and this would have given it an energetic advantage to grow faster than FECA. The endogenized protomitochondrion would have also died faster than the host, based on the well-established linear relationship between growth rate and death rate[27]. Theoretically, this could have led to accumulation of dead protomitochondrion cells in the host cell and, thus, protomitochondrion-derived bacterial ester lipids and bacterial genetic material. Given that ether and ester lipids can mix[28,29] and the host archaeon likely grew slowly, the archaeon's membranes' ether lipids could have been slowly and potentially passively replaced by ester lipids. Moreover, the accumulation of protomitochondrion-derived lipids could have passively increased the host's size. This may help explain (i) why the eukaryote membrane is composed of bacteria-like ester lipids yet is a single layer and lacks a peptidoglycan layer (unlike Gram-negative bacteria and the [proto]mitochondrion) and (ii) how the protoeukaryote cell size may have began to increase. Next, regarding the bacterial genetic material release from protomitochondrion death, such exposure to intracellular foreign genetic material may have made the host archaeon prone to acquisition of bacterial genes—gene transfer from an endosymbiont to the host organism is a common phenomenon. This process may have contributed to high enrichment of bacteria-derived genes in the host genome and perhaps increased evolutionary rates (that is, the endosymbiont may accumulate mutations faster given its higher cell replication rate).

### Supplementary Methods

**Culturing.** During the initial enrichment stage of HC1, we used filter-sterilized brine as the basal medium. The brine was collected in a 5-L glass bottle filled with water from the gas-production well (Extended Data Fig. 1) and immediately sealed with a butyl rubber stopper and screw cap to maintain anoxic conditions. The bottle was brought back to the laboratory under refrigerated conditions in the dark. Upon arrival, the brine was filtered through a 0.22-µm-pore-size polyethersulfone filter unit (Corning) in an anaerobic chamber (Coy Laboratory Products) equipped with a KOACH T 500-F tabletop air filtration system (KOKEN Ltd). Using a 50-ml syringe attached to a 22-gauge needle, 20 ml of the filtered brine was dispensed into sterile anoxic 50-ml serum glass bottles prepared ahead of time (sealed with butyl rubber stoppers and aluminum caps, purged with $N_2/CO_2$ gas (80:20, v/v), and autoclaved). The filtered-sterilized brine was used as a basal medium from the primary to the third subculture. After the fourth subculture, we used an artificial KTG3 medium as the basal medium. The composition of the KTG3 medium was based on the chemical analysis of the brine water[30] (Extended Data Fig. 1) and was as follows (per liter): 1.64 g $MgCl_2 \cdot 6H_2O$, 0.38 g $CaCl_2 \cdot 2H_2O$, 17.64 g NaCl, 0.38 g KCl, 0.5 g $NH_4Cl$, 0.136 g $KH_2PO_4$, 2.3 g $NaHCO_3$, 10 ml $SrCl_2$ solution, 10 ml NaF/LiCl solution, 1 ml trace element solution, 2 ml Se/W solution, 0.5 ml resazurin solution, 1 ml vitamin solution, and 25 ml $Na_2S$/cysteine solution. The composition and preparation of all the solutions were as previously described[31]. The artificial medium was purged with $N_2/CO_2$ gas (80:20 v/v), and the pH was adjusted to 7.0 at 25°C.

The purity of HC1 was routinely examined by microscopy and iTag analysis. Purity was further verified by whole genome shotgun sequencing, which only detected the sequences of HC1 and *Methanocalculus* MC2. The preceding "mother" enrichment cultures of the HC1 and MC2 co-culture, three microbial populations had persisted: *Mariniphaga*, *Alcanivorax*, or *Methanolobus*. Based on separate isolation of these strains, we had confirmed medium compositions they grow under: (i) Marine Broth 2216; (ii) 1/5Marine Broth 2216 plus 0.5% pyruvate; and (iii) KTG3 medium supplemented with 5 mM methanol, respectively (all at 30°C; see below). Using this information, we tested whether we could detect any growth of these organisms in subcultures from the HC1 and MC2 co-culture—that is, ensure that the three above populations had been rejected. In addition, to test for any other contaminating organotrophic organisms, we also incubated subcultures in the following media at 10°C, 20°C, 30°C, 37°C, and 55°C: (i) thioglycolate medium (BD Difco) and (ii) KTG3 medium supplemented with 1 mM sucrose, 1 mM glucose, 1 mM fructose, 1 mM xylose, and 0.01% (w/v) yeast extract. We did not detect any growth of contaminating bacteria/archaea after two months of incubation in all five media. We also used PCR with bacterial 16S rRNA gene primer pairs 27F/907R[32,33] and EUB338*/1490R[32,34–36] and HCR-FISH with 16S rRNA-targeted oligonucleotide probe targeting bacteria EUB338[34] to verify that no contaminating bacteria are present in the HC1 and MC2 co-cultures.

For the cultivation of other *Promethearchaeota* members that may not have been selected for using the enrichment procedure, we used a basal medium that successfully cultured *P. syntrophicum* strain MK-D1 (MK medium)[31]. SC1 was obtained from a culture using MK medium amended with propionate and antibiotics (Supplementary Table 1). After obtaining a stable culture of SC1, we performed HCR-FISH and PCR in the same way as for the purity check of HC1, but no bacterial signals were detected, suggesting that SC1 is a pure tri-culture consisting of only three archaeal species. We also confirmed the absence of microbial growth other than SC1 in the following media at 10°C, 20°C, 30°C, 37°C, and 55°C for 3 months of incubation: (i) thioglycolate medium (BD Difco); and (ii) MK medium supplemented with 1 mM sucrose, 1 mM glucose, 1 mM fructose, 1 mM xylose, and 0.01% (w/v) yeast extract.

To isolate *Mariniphaga* and *Alcanivorax* from the HC1 enrichment culture, we used Marine Broth 2216 and 1/5 Marine Broth 2216 plus 0.5% sodium pyruvate, respectively, and cultured at 30°C. The isolated bacterial strains were used to test their susceptibility to antibiotics (Supplementary Table 2).

**Immunofluorescence staining using lokiactin-specific antibodies.** For western blotting, archaeal cultures (270 ml of HC1, and 80 ml of SC1) were concentrated to about 5 ml using a 0.22-µm-pore-size polyethersulfonate (PES) filter unit (Corning) on a clean bench. The concentrated cell suspensions were further concentrated by centrifugation at 20,400g for 10 min at 4°C. The collected cell samples were washed once with their respective basal media without energy substrates (e.g., peptone and yeast extract) and then denatured in SDS sample buffer containing 1% (v/v) dithiothreitol at 97°C for 5 min. The samples were run on Super Sep Ace 10–20% precast polyacrylamide gel (Fujifilm), transferred to PVDF membranes, and blocked with Blocking One (Nacalai tesque). The blots were probed with respective primary antibodies and secondary antibody (goat anti-rabbit IgG (H+L) HRP conjugate). Signals of the blots were detected by chemiluminescence with ECL Select reagent (Cytiva). The yeast (*Saccharomyces cerevisiae*) lysate used as the negative control was prepared in accordance with a previous report[37].

For immunofluorescence staining, cells were fixed under anaerobic conditions by adding a mixture of formaldehyde (methanol-free, formaldehyde solution, 16% w/v, Electron Microscopy Sciences) and glutaraldehyde (distilled grade, 50% solution, TAAB

Laboratories Equipment Ltd) directly to the fully grown cultures to final concentrations of 4% (w/v) formaldehyde and 0.2% (w/v) glutaraldehyde for 2 h at room temperature in the dark. The fixed cells were then concentrated by centrifugation at 20,400g for 10 min at 4°C and immobilized on poly-L-lysine-coated 8-well chamber slides. The slide samples were subsequently permeabilized with 0.1% Triton X-100 and blocked with 3% BSA. After blocking, the cells were probed with respective primary antibodies (1:500 diluted) and secondary antibodies diluted either 1:100 (goat anti-rabbit Alexa Fluor 555, Abcam) for observation under a BX53 Olympus epifluorescence microscope or 1: 200 (goat anti-rabbit abberior STAR 580, Abberior) for observation with a STELLARIS STED confocal microscope (Leica). The samples were counterstained with SYBR Green-I (Thermo Fisher Scientific) and mounted with Prolong Diamond (Thermo Fisher Scientific) prior to observation. Images were acquired using a STELLARIS STED confocal microscope (Leica) with a ×100/1.4 NA oil-immersion objective at 6 × zoom, line scanning frequency of 400 Hz, and line average of 3. Fluorescence from the secondary antibodies was acquired using stimulated emission depletion (STED) mode with 587 nm for excitation and the depletion at 775 nm, while SYBR Green-I-derived fluorescence was acquired in confocal mode. Through all immunofluorescence staining procedures, pipetting was performed gently to minimize physical stress on the cells and prevent the detachment of cellular protrusions. Although SPY505-DNA (Spirochrome) was initially considered for DNA staining, we opted not to use it because it also stained well non-cell materials such as medium-derived precipitates.

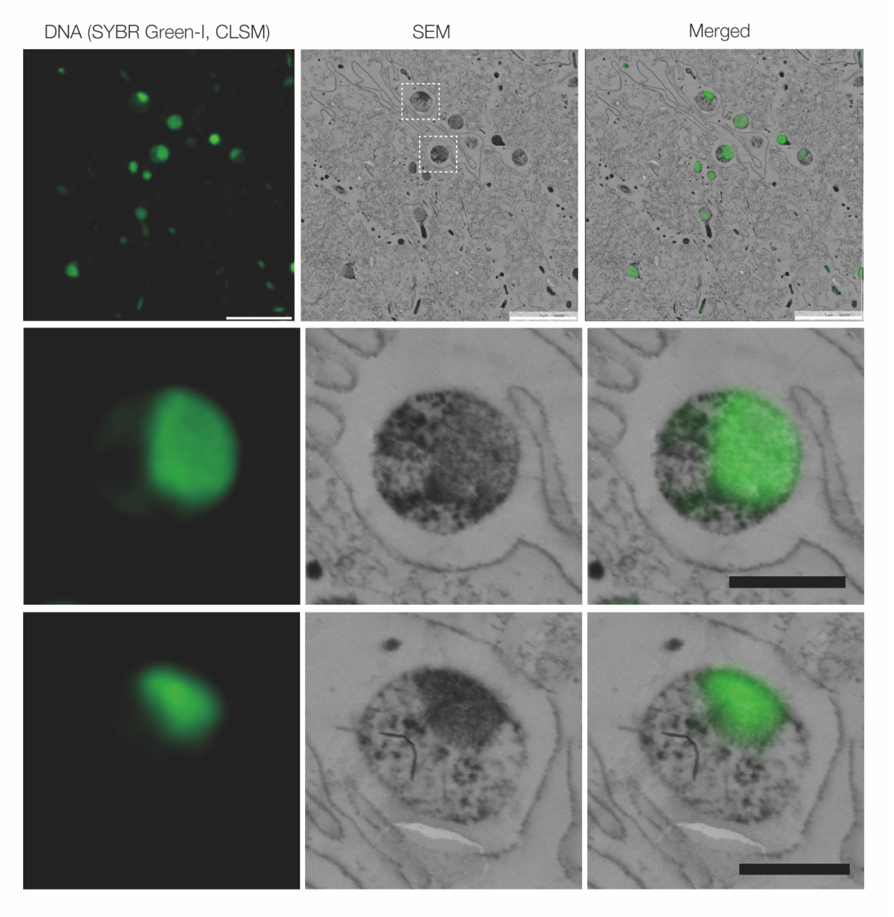


**Supplementary Fig. 1 | CLEM images of HC1.** The lower two panels are enlarged images of the area enclosed by the white dotted-line squares in the top SEM image. Scale bars, 5 μm for the top panel and 1 μm for the lower two panels.

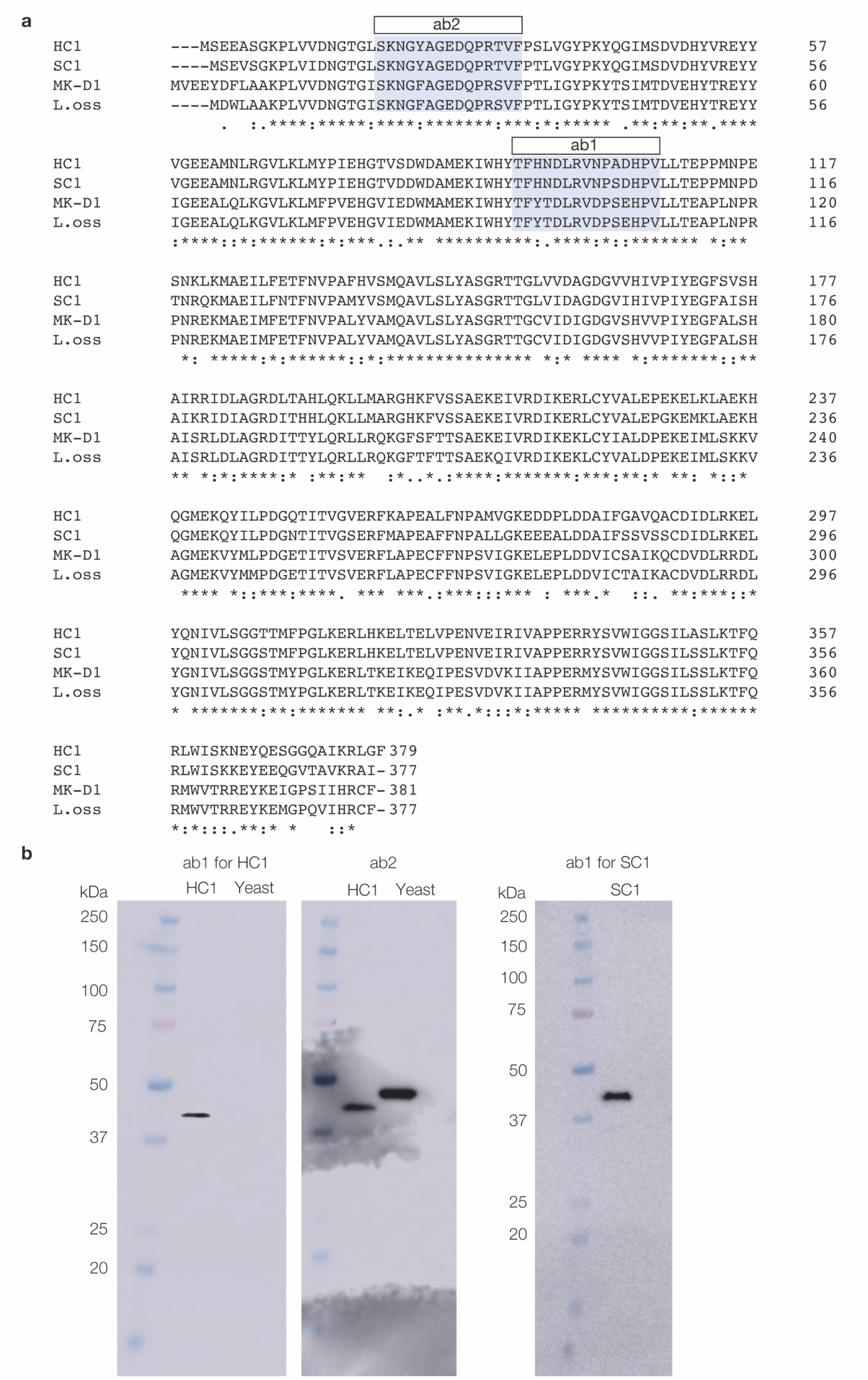


**Supplementary Fig. 2 | Lokiactin amino acid sequence of cultured archaea and confirmation of their expression using Western blotting analysis. a**, Multiple sequence alignment for lokiactin sequences of cultured *Promethearchaeota* strains.

Sequences were aligned with Clustal Omega provided EMBL-EBI as a web service (https://www.ebi.ac.uk/jdispatcher/msa/clustalo). The regions colored in light blue showed that the peptides were used for creating antibodies (*i.e.*, ab1 for HC1, ab1 for SC1, and ab2 for both strains). **b**, Western blotting analysis of lokiactin-specific antibodies. Yeast lysate was used as a negative control during the western blotting analysis using HC1 cultures.

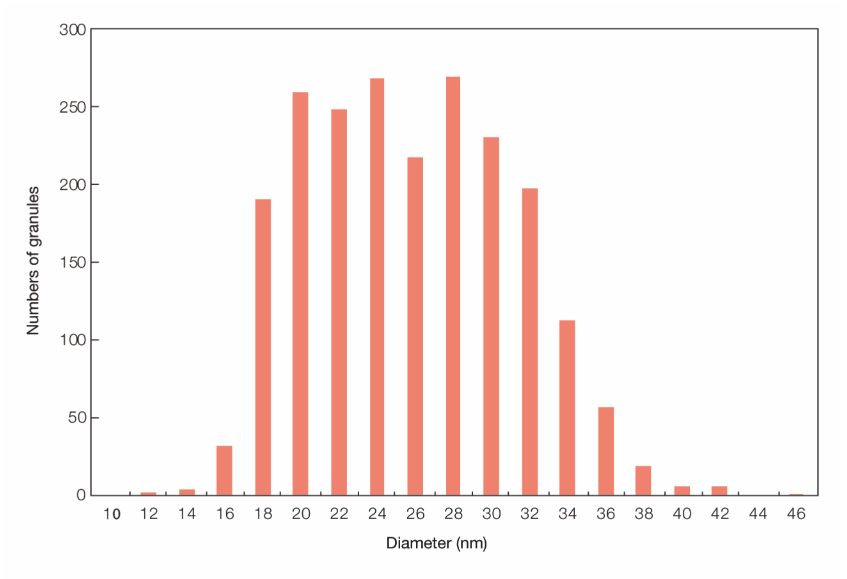


**Supplementary Fig. 3 | Size distribution of granular structures in an HC1 cell.** A total of 2,127 granular structures depicted in Fig. 2l were quantified using AMIRA 3D Pro software.

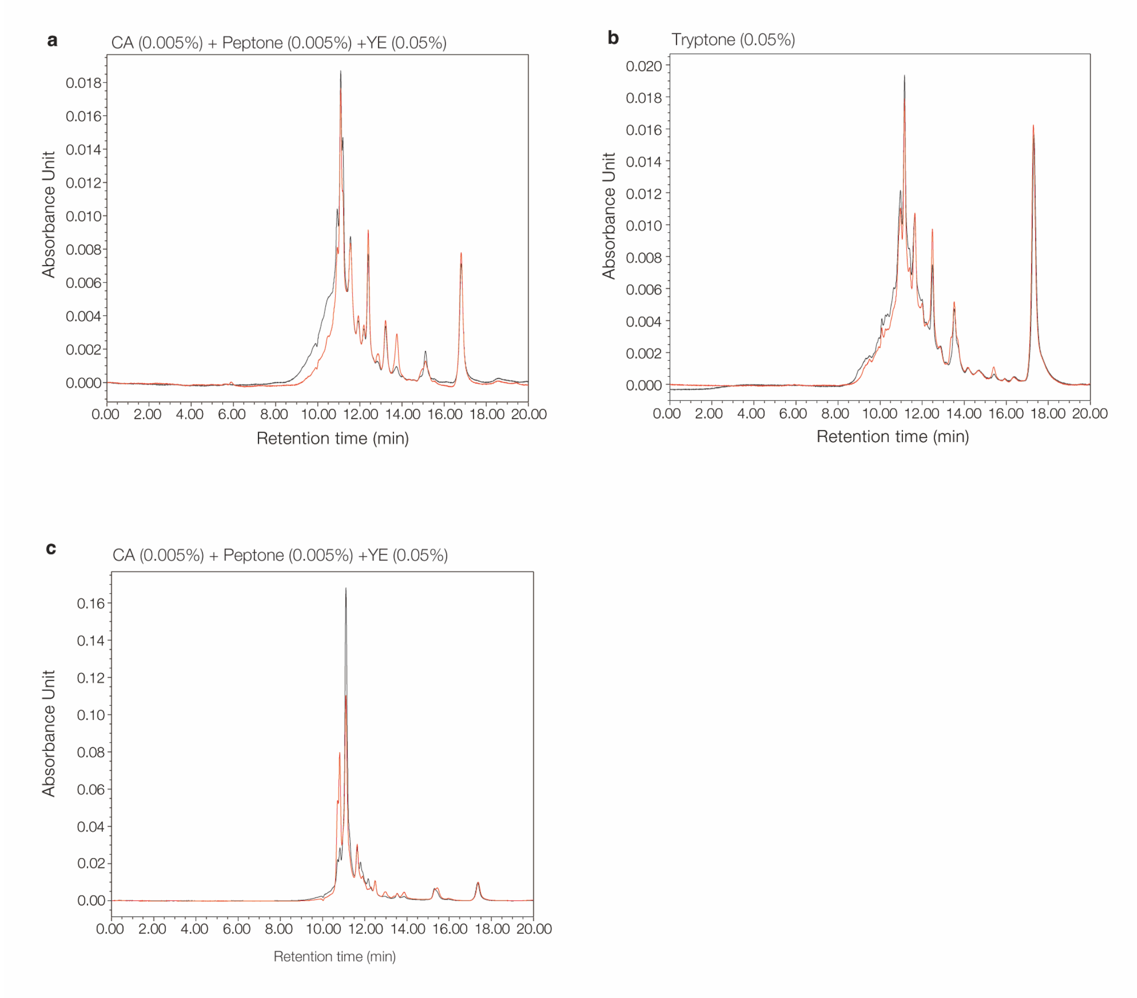


**Supplementary Fig. 4 | UPLC-based detection of high molecular weight compounds in HC1 and SC1 cultures. a**,**b**, HC1 cultures fed with casamino acids, peptone, and yeast extract (0.05%, w/v) (a), and tryptone (0.05%). **c**, SC1 culture fed with casamino acids, peptone and yeast extract (0.005%, 0.005%, and 0.05%, respectively). Black and red lines show before and after cultivation, respectively.

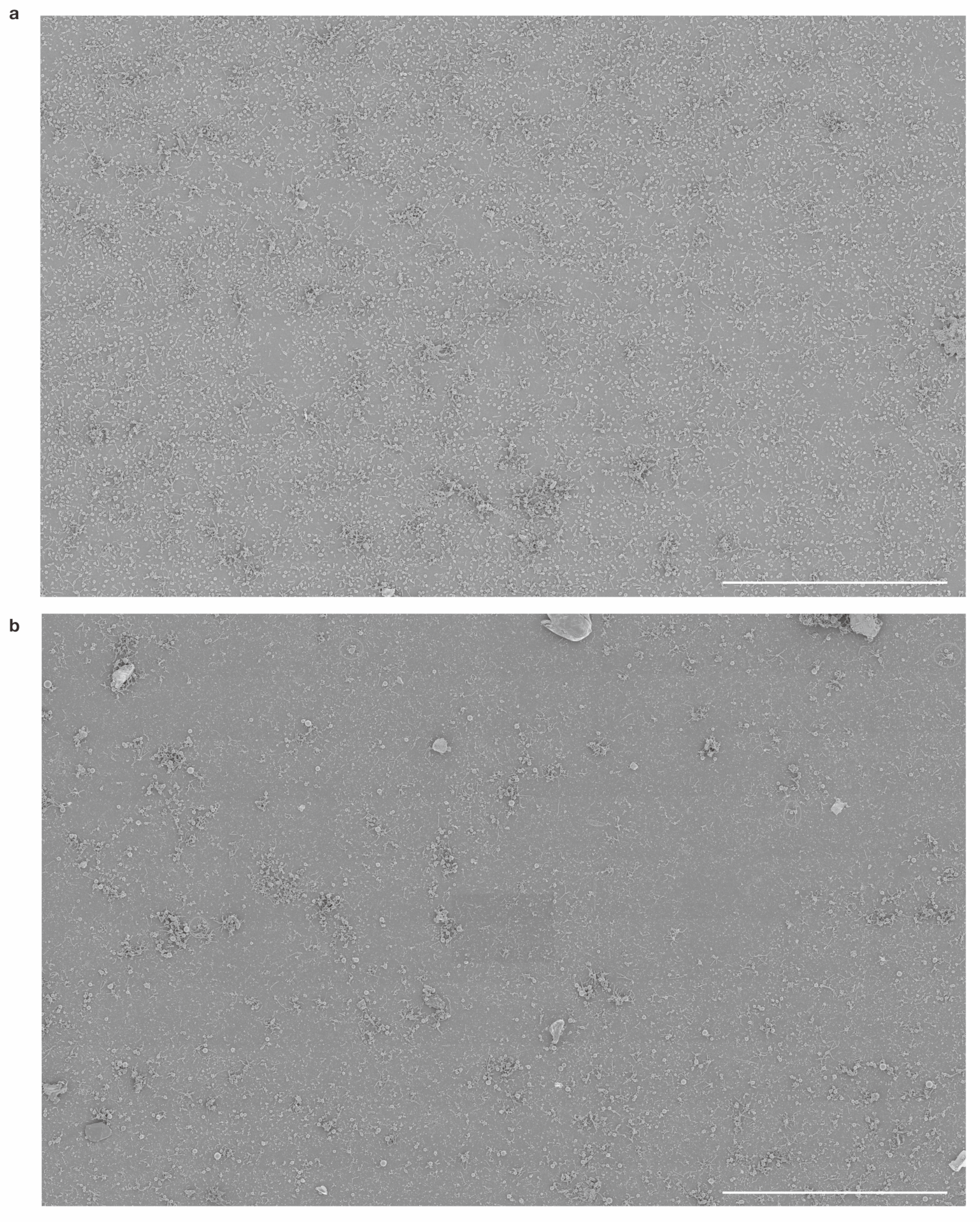


**Supplementary Fig. 5 | Wide-area SEM images of HC1. a, b,** HC1 cultured with 0.005% yeast extract (**a**) and 0.05% yeast extract (**b**). The culture media also contained 0.005% casamino acids and 0.005% peptone. Scale bars, 80μm. High-resolution versions of these images and segmentation images, and data set have been deposited on figshare (https://doi.org/0.6084/m9.figshare.28379207).

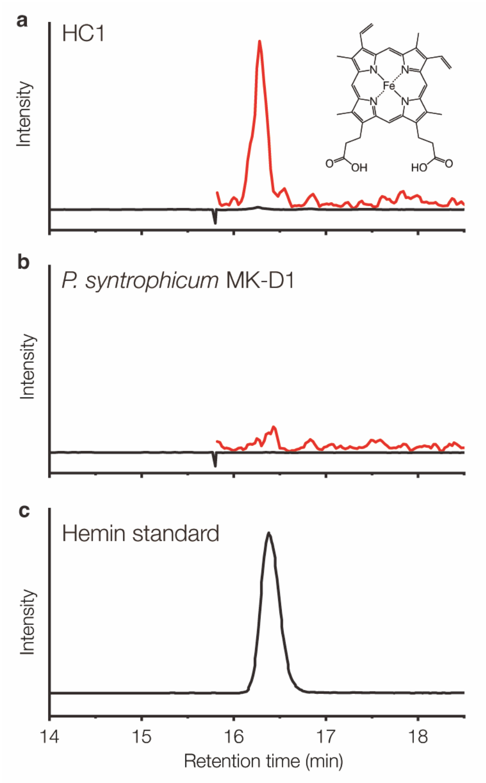


**Supplementary Fig. 6 | Chromatograms of heme B. a**, HC1. **b**, *P. syntrophicum* MK-D1. **c,** Standard material of heme B in oxidized form (hemin; ≥97.0% purity, Sigma-Aldrich). The y-axes for (**a**) and (**b**) are shown on the same scale. Each measurement was performed in triplicate, yielding consistent results across all samples.

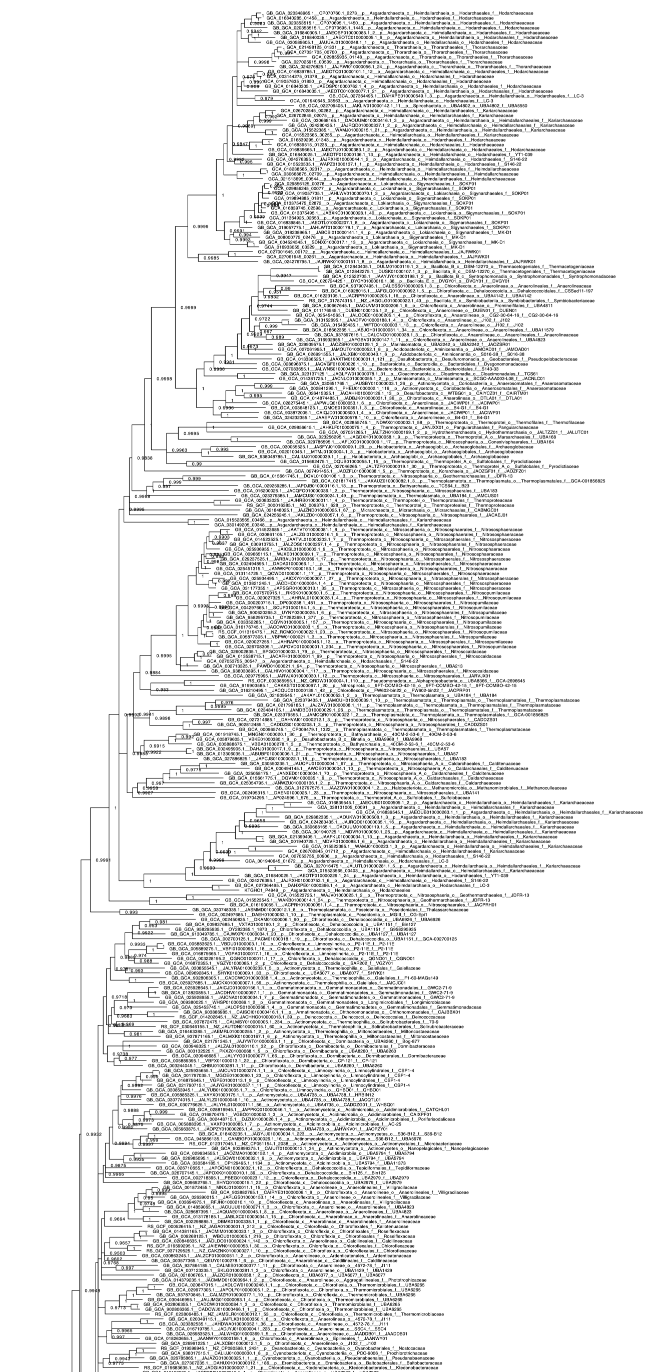

**Supplementary Fig. 7 | Maximum likelihood estimation for the phylogeny of heme O synthase, depicting the ungrouped version of Fig. 4b.** Branches with low ultrafast bootstrap support (<0.95) were collapsed.

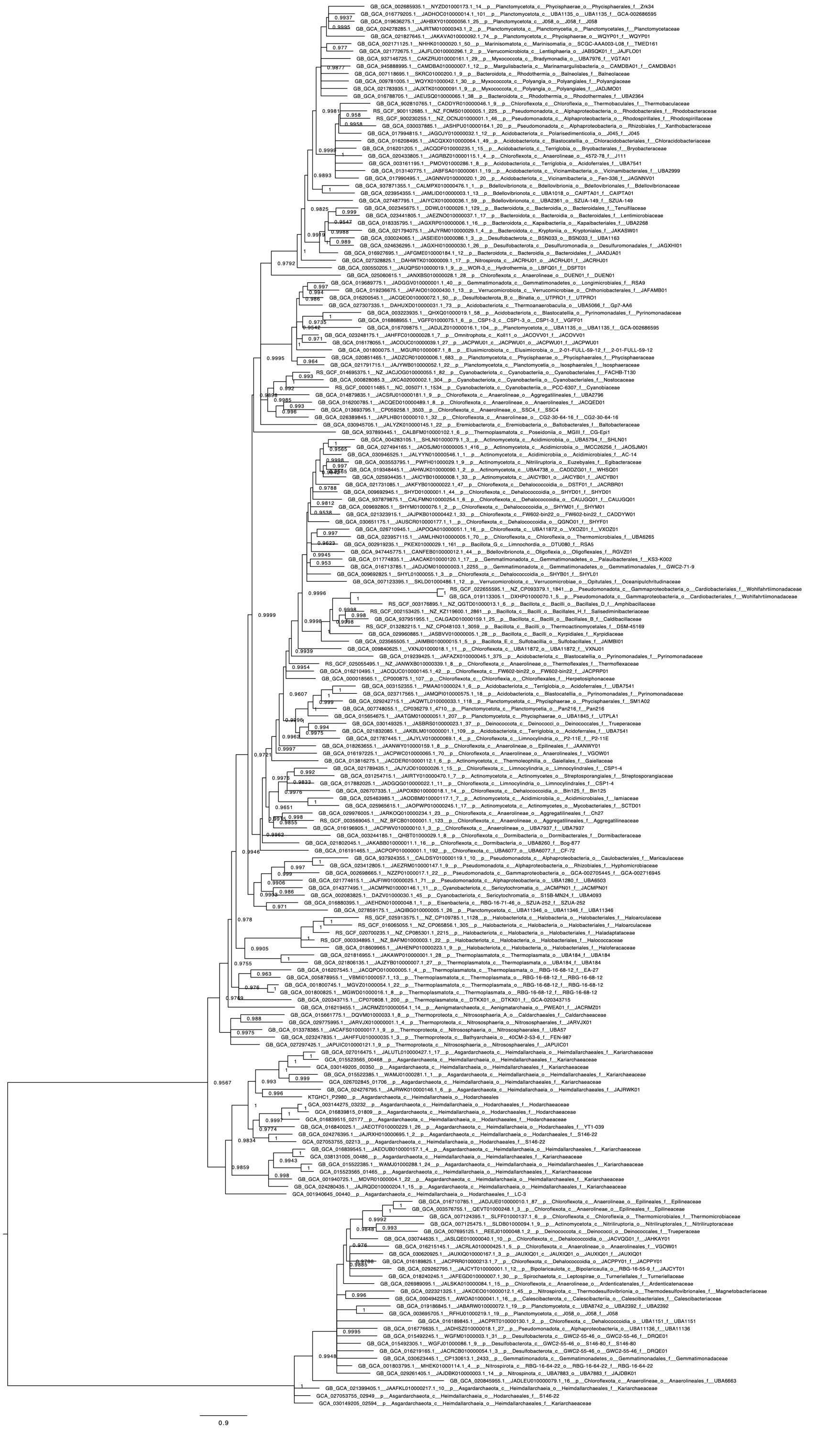

**Supplementary Fig. 8 | Maximum likelihood estimation for the phylogeny of cytochrome *c* oxidase subunit 1, depicting the ungrouped version of Fig. 4c.** Branches with low ultrafast bootstrap support (<0.95) were collapsed.

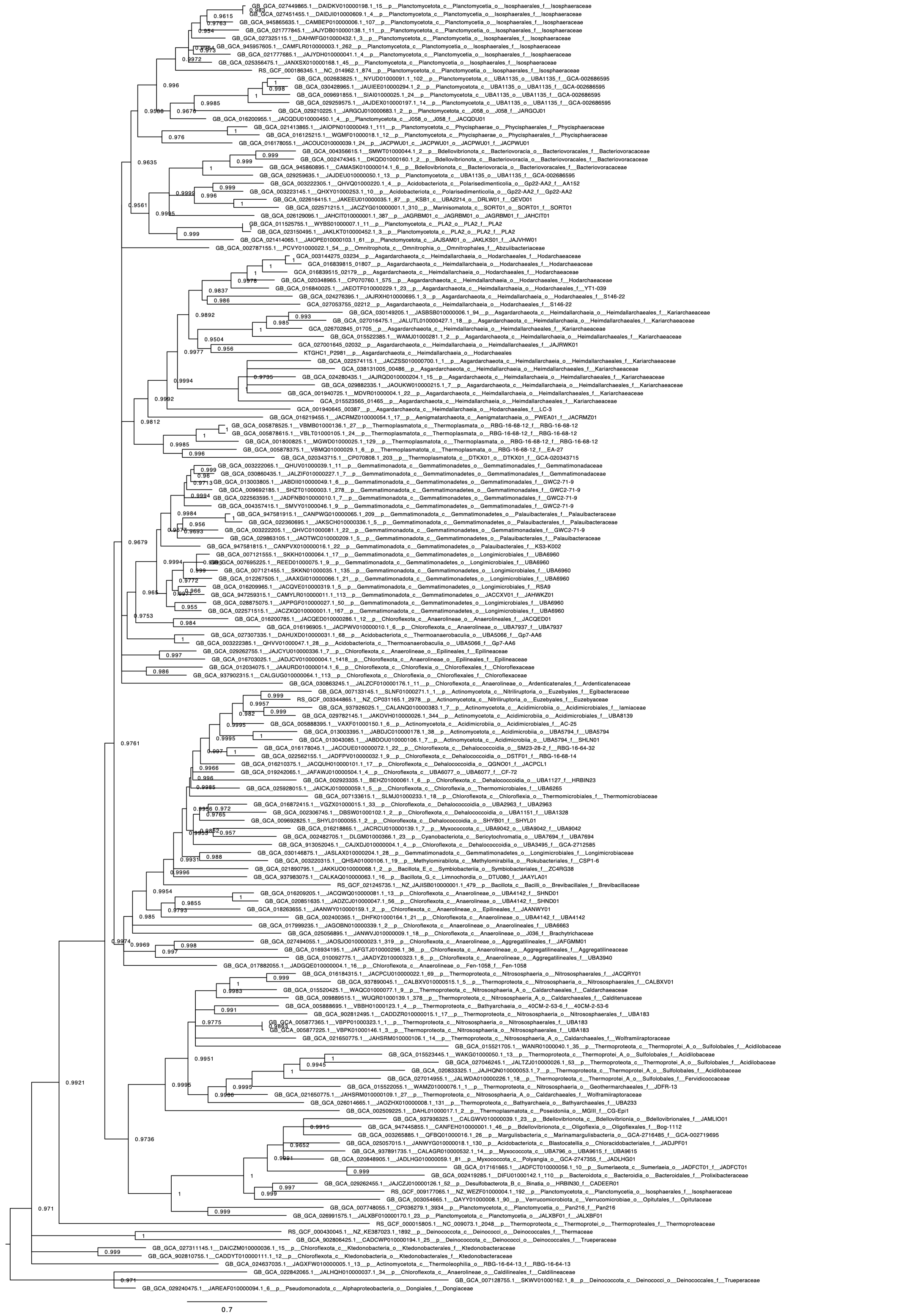

**Supplementary Fig. 9 | Maximum likelihood estimation for the phylogeny of cytochrome *c* oxidase subunit 3, depicting the ungrouped version of Fig. 4d.** Branches with low ultrafast bootstrap support (<0.95) were collapsed.

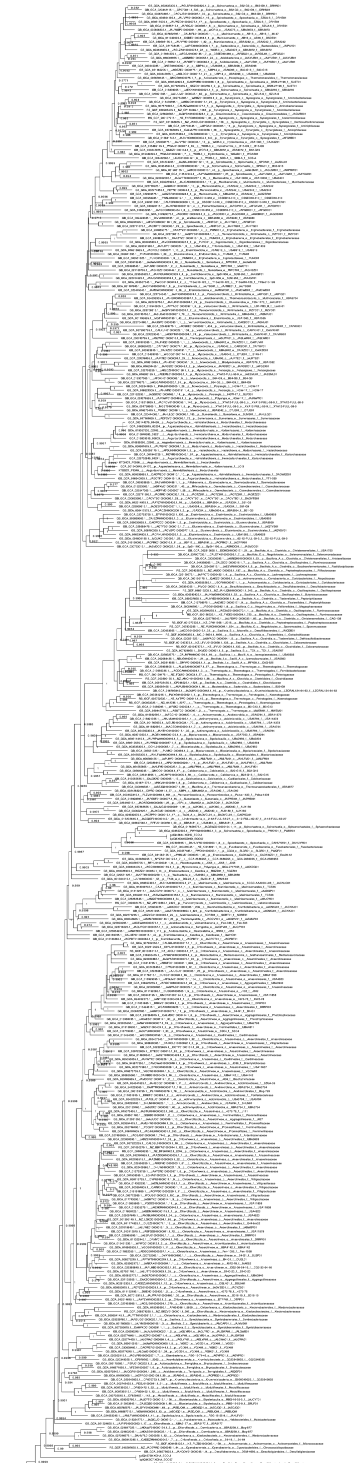

**Supplementary Fig. 10 | Maximum likelihood estimation for the phylogeny of xanthine dehydrogenase, depicting the ungrouped version of Fig. 4e.** Branches with low ultrafast bootstrap support (<0.95) were collapsed.

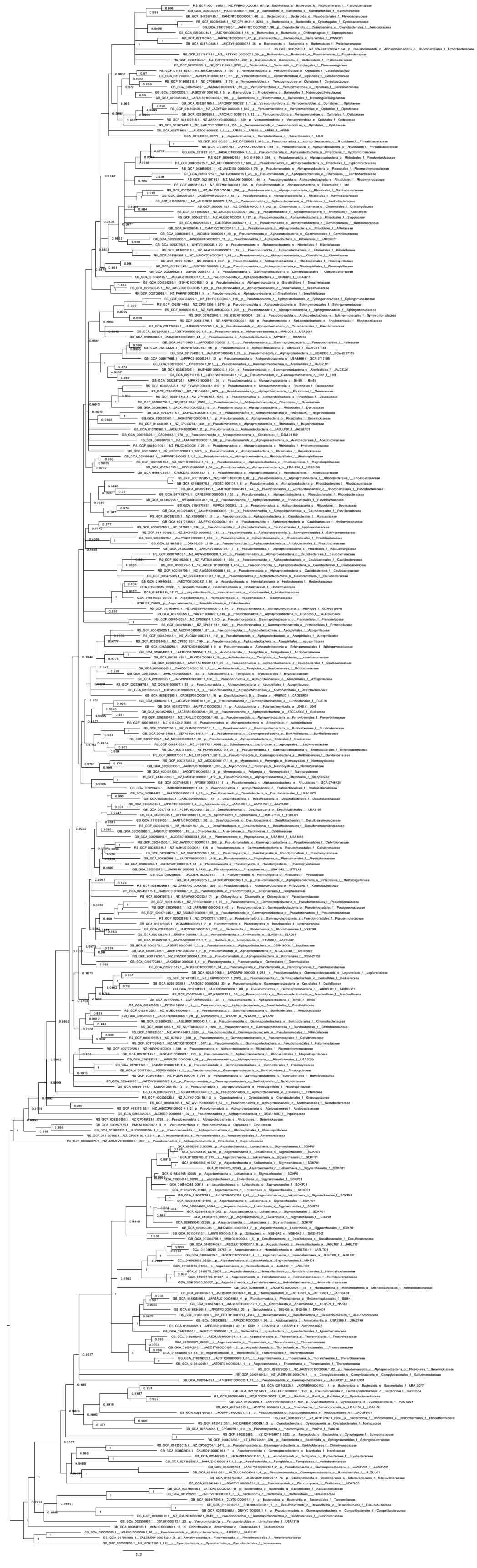

**Supplementary Fig. 11 | Maximum likelihood estimation for the phylogeny of catalase, depicting the ungrouped version of Fig. 4f.** Branches with low ultrafast bootstrap support (<0.95) were collapsed.

### Supplementary references

1. Aoki, M. *et al.* A long-term cultivation of an anaerobic methane-oxidizing microbial community from deep-sea methane-seep sediment using a continuous-flow bioreactor. *PLoS ONE* **9**, e105356 (2014).

2. Imachi, H. *et al.* Cultivation of methanogenic community from subseafloor sediments using a continuous-flow bioreactor. *ISME J.* **5**, 1913–1925 (2011).

3. Girguis, P. R., Orphan, V. J., Hallam, S. J. & DeLong, E. F. Growth and methane oxidation rates of anaerobic methanotrophic archaea in a continuous-flow bioreactor. *Appl. Environ. Microbiol.* **69**, 5472–5482 (2003).

4. Zhu, Q.-Z., Wegener, G., Hinrichs, K.-U. & Elvert, M. Activity of ancillary heterotrophic community members in anaerobic methane-oxidizing cultures. *Front. Microbiol.* **13**, 912299 (2022).

5. Imachi, H. *et al.* Isolation of an archaeon at the prokaryote-eukaryote interface. *Nature* **577**, 519–525 (2020).

6. Imachi, H. *et al.* *Promethearchaeum syntrophicum* gen. nov., sp. nov., an anaerobic, obligately syntrophic archaeon, the first isolate of the lineage 'Asgard' archaea, and proposal of the new archaeal phylum *Promethearchaeota* phyl. nov. and kingdom *Promethearchaeati* regn. nov. *Int. J. Syst. Evol. Microbiol.* **74**, 006435 (2024).

7. Janssen, P. H. Selective enrichment and purification of cultures of *Methanosaeta* spp. *J. Microbiol. Meth.* **52**, 239–244 (2003).

8. Martens-Habbena, W., Berube, P. M., Urakawa, H., Torre, J. R. de la & Stahl, D. A. Ammonia oxidation kinetics determine niche separation of nitrifying Archaea and Bacteria. *Nature* **461**, 976–979 (2009).

9. Russum, S. *et al.* Comparative population genomic analyses of transporters within the Asgard archaeal superphylum. *PLoS ONE* **16**, e0247806 (2021).

10. Liu, L., Li, S., Han, J., Lin, W. & Luo, J. A two-step strategy for the rapid enrichment of *Nitrosocosmicus*-like ammonia-oxidizing *Thaumarchaea*. *Front. Microbiol.* **10**, 875 (2019).

11. Bayer, B. *et al.* Physiological and genomic characterization of two novel marine thaumarchaeal strains indicates niche differentiation. *ISME J.* **10**, 1051–1063 (2016).

12. Simon, H. M. *et al.* Cultivation of mesophilic soil crenarchaeotes in enrichment cultures from plant roots. *Appl. Environ. Microbiol.* **71**, 4751–4760 (2005).

13. Dridi, B., Raoult, D. & Drancourt, M. Archaea as emerging organisms in complex human microbiomes. *Anaerobe* **17**, 56–63 (2011).

14. Stieglmeier, M. *et al. Nitrososphaera viennensis* gen. nov., sp. nov., an aerobic and mesophilic, ammonia-oxidizing archaeon from soil and a member of the archaeal phylum *Thaumarchaeota*. *Int. J. Syst. Evol. Microbiol.* **64**, 2738–2752 (2014).

15. Thombre, R. S., Shinde, V., Thaiparambil, E., Zende, S. & Mehta, S. Antimicrobial activity and mechanism of inhibition of silver nanoparticles against extreme halophilic archaea. *Front. Microbiol.* **7**, 1424 (2016).

16. Spang, A. *et al.* Proposal of the reverse flow model for the origin of the eukaryotic cell based on comparative analyses of Asgard archaeal metabolism. *Nat. Microbiol.* **4**, 1138–1148 (2019).

17. Eme, L. *et al.* Inference and reconstruction of the heimdallarchaeial ancestry of eukaryotes. *Nature* **618**, 992–999 (2023).

18. Bulzu, P.-A. *et al.* Casting light on Asgardarchaeota metabolism in a sunlit microoxic niche. *Nat. Microbiol.* **4**, 1129–1137 (2019).

19. Liu, Y. *et al.* Expanded diversity of Asgard archaea and their relationships with eukaryotes. *Nature* **593**, 553–557 (2021).

20. Appler, K. E. *et al.* Oxygen metabolism in descendants of the archaeal-eukaryotic ancestor. *Preprint at https://www.biorxiv.org/content/10.1101/2024.07.04.601786v1* (2024).

21. Nakagawa, S. *et al.* Characterization of protein glycosylation in an Asgard archaeon. *BBA Adv.* **6**, 100118 (2024).

22. Nakagawa, S. *et al.* N-linked protein glycosylation in *Nanobdellati* (formerly DPANN) archaea and their hosts. *J. Bacteriol.* **206**, e00205-24 (2024).

23. Jarrell, K. F. *et al.* N-linked glycosylation in *Archaea*: a structural, functional, and genetic analysis. *Microbiol. Mol. Biol. Rev.* **78**, 304–341 (2014).

24. Fujinami, D., Taguchi, Y. & Kohda, D. Asn-linked oligosaccharide chain of a crenarchaeon, Pyrobaculum calidifontis, is reminiscent of the eukaryotic high-mannose-type glycan. *Glycobiology* **27**, 701–712 (2017).

25. Villanueva, L. *et al.* Bridging the membrane lipid divide: bacteria of the FCB group superphylum have the potential to synthesize archaeal ether lipids. *ISME J.* **15**, 168–182 (2020).

26. Villanueva, L., Schouten, S. & Damsté, J. S. S. Phylogenomic analysis of lipid biosynthetic genes of Archaea shed light on the "lipid divide". *Environ. Microbiol.* **19**, 54–69 (2017).

27. Biselli, E., Schink, S. J. & Gerland, U. Slower growth of *Escherichia coli* leads to longer survival in carbon starvation due to a decrease in the maintenance rate. *Mol. Syst. Biol.* **16**, MSB209478 (2020).

28. Caforio, A. *et al.* Converting *Escherichia coli* into an archaebacterium with a hybrid heterochiral membrane. *Proc. Natl. Acad. Sci. USA* **115**, 3704–3709 (2018).

29. Hoekzema, M., Jiang, J. & Driessen, A. J. M. Optimizing archaeal lipid biosynthesis in *Escherichia coli*. *ACS Synth. Biol.* **13**, 2470–2479 (2024).

30. Urai, A. *et al.* Origin of deep methane associated with a unique community of microorganisms in an organic- and iodine-rich aquifer. *ACS Earth Space Chem.* **5**, 1–11 (2021).

31. Imachi, H. *et al.* Cultivation of previously uncultured microorganisms with a continuous-flow down-flow hanging sponge (DHS) bioreactor, using a syntrophic archaeon culture obtained from deep marine sediment as a case study. *Nat. Protoc.* **17**, 2784–2814 (2022).

32. Weisburg, W. G., Barns, S. M., Pelletier, D. A. & Lane, D. J. 16S ribosomal DNA amplification for phylogenetic study. *J. Bacteriol.* **173**, 697–703 (1991).

33. Lane, D. J. 16S/23S rRNA sequencing. in *Nucleic Acid Techniques in Bacterial Systematics* (eds. Stackebrandt, E. & Goodfellow, M.) 115–175 (John Wiley & Sons, 1991).

34. Amann, R. I. *et al.* Combination of 16S rRNA-targeted oligonucleotide probes with flow cytometry for analyzing mixed microbial populations. *Appl. Environ. Microbiol.* **56**, 1919–1925 (1990).

35. Daims, H., Brühl, A., Amann, R., Schleifer, K. H. & Wagner, M. The domain-specific probe EUB338 is insufficient for the detection of all Bacteria: development and evaluation of a more comprehensive probe set. *Syst. Appl. Microbiol.* **22**, 434–444 (1999).

36. Hatamoto, M., Imachi, H., Yashiro, Y., Ohashi, A. & Harada, H. Diversity of anaerobic microorganisms involved in long-chain fatty acid degradation in methanogenic sludges as revealed by RNA-based stable isotope probing. *Appl. Environ. Microbiol.* **73**, 4119–4127 (2007).

37. Kushnirov, V. V. Rapid and reliable protein extraction from yeast. *Yeast* **16**, 857–860 (2000).
